# Pregnancy History and Loss of Normal PSEN2 Function Impact Female Late-Life Cognitive Processing, Inhibitory Signaling, and Gliosis

**DOI:** 10.64898/2026.10.28.762624

**Authors:** Jennie I. Fang, Leanne Lehmann, Katherine E. Prater, Melissa Barker-Haliski

**Affiliations:** Department of Pharmaceutics, University of Washington; Neurosciences Graduate Program, University of California, San Diego; Department of Neurology, University of Washington

**Author notes:** Corresponding Author, Department of Pharmaceutics, University of Washington, Seattle, WA USA.

**Keywords:** Brain-derived neurotrophic factor, Astrocytes, Oligodendrocytes, Parvalbumin, Hippocampal learning and memory, Anxiety-like behavior

## Abstract

**Background:** Alzheimer’s disease (AD) is well-documented to be more progressive in affected females. It is thus critical to define how female-specific life experiences interact with AD-related risk factors to impact disease trajectory and functional burden. For example, pregnancy exerts dramatic influences on the maternal immune system, behavior, and neuroplasticity, specifically in hippocampal and limbic structures. The long-term clinical impacts from pregnancy history (parity) on late-life AD outcomes are, however, conflicting. Thus, we investigated the late-life functional impacts of parity on rodent cognitive function, inhibitory neuron density, and glial biology with loss of normal function of the AD-related risk factor, presenilin 2 (PSEN2).

**Methods:** Female PSEN2 knockout (KO) and wild-type (WT) mice with (multiparous) and without (nulliparous) pregnancy history were challenged at 10-15 months-old in the T-maze (TM) and Open Field Test (OFT), followed by immunofluorescence for neuronal signaling and gliosis markers.

**Results:** Neither parity history nor PSEN2 deficiency impacted hippocampally-mediated spatial working memory of aged female mice in the TM. However, both PSEN2 KO and multiparous mice exhibited decreased OFT locomotor behavior. Multiparous mice also showed reduced OFT center zone exploration (p=0.0045) suggesting a more anxious phenotype. Parity history generally increased inhibitory marker expression in dorsal hippocampus (p=0.017), prefrontal cortex (p=0.021), and amygdala (p=0.035). Parity generally blunted oligodendrocyte marker expression in CA1 (p<0.05), while boosting astrocyte and microglia marker expression in CA3 of dorsal hippocampus.

**Conclusion:** This work suggests that although parity history and PSEN2 deficiency exert individual differing age-related effects on cognition and neurobiology, they do not act synergistically.

## INTRODUCTION

Advanced age and female sex are two major non-modifiable risk factors for Alzheimer’s disease (AD). AD is not only the most well-known age-related neurodegenerative disorder but also exhibits a well-documented sex burden disparity (“2025 Alzheimer’s Disease Facts and Figures,” 2025). Females demonstrate greater prevalence, more rapid cognitive decline following the mild cognitive impairment stage, worsened neuropathological progression, and more sex-specific AD-related risk factors (Emrani & Sundermann, 2025; Pike, 2017). Despite the rapidly aging worldwide population increasing AD prevalence, the age-related changes that underlie sex-specific differences in cognitive burden of this disease are poorly understood. To better understand the cognitive burden differences between males and females with AD, it is crucial to examine how female-specific life experiences, such as pregnancy history, additively or synergistically interact with AD-associated risk factors to impact late life cognitive function.

AD is the most common cause of dementia, accounting for 60-80% of cases, and is characterized by progressive memory loss, executive function decline, and behavioral changes (“2025 Alzheimer’s Disease Facts and Figures,” 2025; Sheppard & Coleman, 2020). Synapse loss and neuronal atrophy occur in the hippocampus and cerebral cortex, and the hallmark amyloid-beta and hyperphosphorylated tau protein aggregates accumulate broadly throughout various brain regions.

While most AD cases are late onset, roughly 5-10% of cases manifest before age 65 and are thus classified as early onset AD (EOAD). Three autosomal dominant risk genes lead to EOAD, including amyloid precursor protein (APP) duplication events and gene variants encoding the homologous proteins, Presenilin 1 and 2 (PSENs 1 and 2). PSEN2 variants are the least common genetic cause of EOAD, however, this could potentially be due to the later age of symptom onset (∼40-70 year-old), which may lead to PSEN2 variant-related AD being masked as late-onset AD (Blauwendraat et al., 2016; Course et al., 2022). Furthermore, when compared with other AD-associated risk genes, PSEN2 is relatively understudied and insufficiently characterized, necessitating increased research on how PSEN2 variants contribute to AD pathophysiology. Like late-onset AD, EOAD exhibits sex differences.

While EOAD prevalence does not differ between males and females, there is greater cognitive impairment and broader brain atrophy in females versus males, when compared to sex-matched unaffected individuals (Contador et al., 2022; Peeters et al., 2022). Thus, despite the largely genetic causes of EOAD, the more common late-onset AD shares conserved clinical and pathological features when the influence of biological sex is considered.

PSEN2 is a γ-secretase subunit that cleaves APP into the ß-amyloid peptide form that typically aggregates into the characteristic plaques of AD (Cai et al., 2015). The most common PSEN2 gene variants lead to a loss of normal γ-secretase function such that deletion of PSEN2 gene expression (i.e., PSEN2 knockout – KO) results in a similar biochemical outcome *in vivo* (Jayadev et al, 2010; 2013). Thus, PSEN2 KO models are suitable to *a priori* define how loss of normal PSEN2 expression impacts long-term physiological processes in normal aging and disease. Despite this, PSEN2 KO mice do not exhibit significantly altered APP metabolism or ß-amyloid plaque deposition with advanced age (Herreman et al., 1999a; Jayadev et al., 2010; Robinson-Cooper et al., 2026), suggesting that loss of normal PSEN2 function may also underlie AD onset in an amyloid-independent manner (Herreman et al., 1999; Jayadev et al., 2010). Instead, AD-associated PSEN2 variants show increased microglia reactivity and heightened inflammatory responses (Jayadev et al., 2010, 2013; Luquez et al., 2026).

Other non-amyloidogenic functions of normal PSEN2 function involve modulation of the Wnt/ß-catenin signaling pathway, cellular Ca^2+^ homeostasis regulation, transcriptional regulation via Notch signaling, and cleavage of voltage-gated sodium channels (Jurisch-Yaksi et al., 2013; Pizzo et al., 2020; Kim et al., 2005), all pathways and processes that can be disrupted with normal aging or comorbid conditions. For example, we have reported that PSEN2 KO mice exhibit differential sex-based cognitive deficits with experimentally-induced neuronal hyperexcitability (Knox et al., 2023), suggesting that age-related pathological impacts of PSEN2 deficiency may uniquely affect females. During pregnancy, the maternal immune system is modulated to tolerate the half-foreign fetus while still retaining capacity to combat pathogens, a feat that heavily involves microglia as the brain-resident immune cells (Gumusoglu & Stelzer, 2025). Microglia number, morphology, and dynamics are vastly altered in the postpartum brain (Rim et al., 2026). Thus, the PSEN2 KO mouse provides an interesting platform to study how the transient changes in immunological function during pregnancy interact with AD-associated risk factors to impact late-life neurobiology, neuroinflammation, and cognitive function.

Pregnancy history (i.e., parity) confers conflicting effects on the development and risk for clinical AD diagnosis and burden in humans. The number of pregnancies correlates with severity of clinical AD neuropathological lesions (Beeri et al., 2009), decreased age of AD onset (Sobow & Kloszewska, 2003), and increased AD risk (Colucci et al., 2006; Jang et al., 2018), suggesting that pregnancy impacts long-term AD burden. However, others have reported that there is no interaction between parity and later AD development in females. For example, there is an inverted U-shaped association between parity and dementia risk in women and men (Basit et al., 2023; Gong et al., 2022), indicating that sociocultural, rather than biological, factors influence female outcomes and mediate any AD- pregnancy interaction. Conversely, research supporting neuroprotective effects of parity also exists; women with more cumulative months pregnant have lower AD risk (M. Fox et al., 2018). A potential explanation for these contrasting findings is that societal and behavioral factors, such as variation in child rearing and family life, may confound clinical studies (Beeri et al., 2009). Yet, rodent investigations addressing this interaction are limited. For example, although wild-type (WT) non-breeder females outperform wild-type breeders in various memory parameters, AD-associated breeders do not show improved cognition versus genotype-matched non-breeders (Cui et al., 2014). In a rat AD model (Lee et al., 2024), spatial memory task performance of parous rodents involved search strategies that were largely reflective of decreased hippocampal activation. Altogether, the interaction between parity and AD-associated genetic risk factors on long-term functional and cognitive outcomes remains unclear.

During pregnancy, not only does the maternal immune system adapt, but the maternal brain also undergoes significant changes, altering cognition (Duarte-Guterman et al., 2023; Eid et al., 2019; Trainer et al., 2006). Cellular neuroplasticity, neurogenesis, synaptic remodeling, and astrocytic density changes in the maternal brain (Barba-Müller et al., 2019). Additionally, estrogen and progesterone level fluctuations dramatically modify the GABA (γ-aminobutyric acid) and glutamate-mediated inhibition and excitation (Gilfarb & Leuner, 2022; Zhao & Gammie, 2014). However, the mechanisms that underlie parity-related changes in memory and behavior across the lifespan in individuals with AD-related genetic risk factors is understudied. We therefore hypothesized that parity history and loss of normal PSEN2 function could alter behavioral outcomes, as well as limbic system and prefrontal cortex neurobiology and neuroinflammation with advanced age.

## METHODS

### Animals

Female 10-15-month-old PSEN2 KO mice were bred at the University of Washington (UW) from stock originally acquired from the Jackson Laboratory (stock #005617), as previously reported (Beckman et al., 2020). All breeding at UW was between PCR-confirmed PSEN2 KO pairs. Age-matched female WT (stock #000664) were acquired from Jackson Labs at 12 weeks-old and aged alongside KO mice until behavioral testing 7-12 months later. Multiparous WT and KO females were retired breeders, with experimental testing conducted at minimum 7 weeks after the last litter, with the number of pregnancies/mouse unknown aside from confirming parity history. We did not track estrous cycles.

Animals were housed on a 14:10 light cycle (lights on a 06 h00: off at 20 h00) in ventilated cages with corncob bedding in a manner consistent with the *Guide for the Care and Use of Laboratory Animals* (protocol 4387-01) and all animal work was approved by the UW Institutional Animal Care and Use Committee and conformed to ARRIVE guidelines (Sert et al., 2020). Mice had free access to irradiated chow (PicoLab Rodent Diet 20–5053), water, and enrichment, except during periods of behavioral testing. Multiparous females received a high-fat breeder chow (PicoLab Rodent Diet 20–5058) from breeder status initiation (7-10 weeks) until retirement (∼10 months). While both formulas contain roughly 20% protein, 5058 is a high-energy diet formulated with double the fat content (9.0%) compared to 5053 (4.5%) to support demanding mouse breeding in transgenic strains. Behavioral testing was performed between 10h00 and 17h00 beginning at 11-13-months-old (Figure 1A).

**Figure 1.**
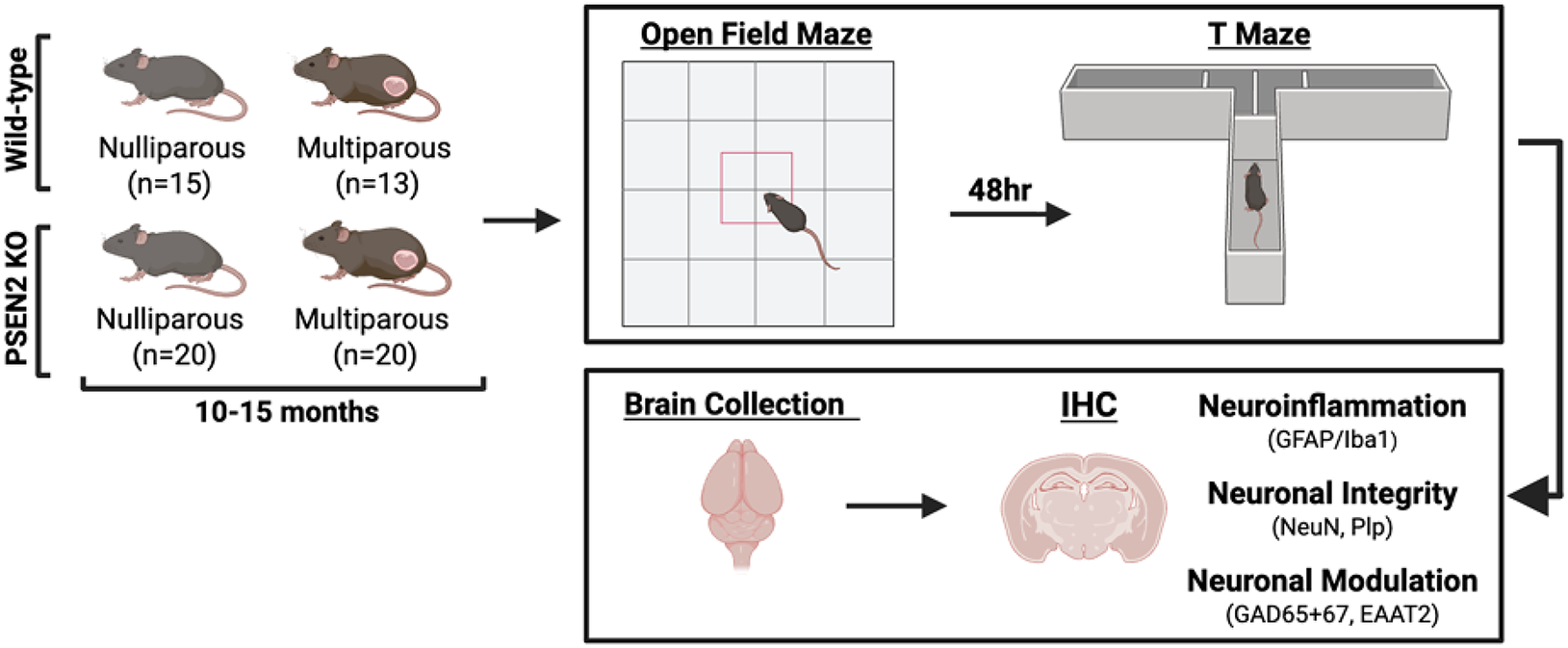
Experimental design to assess the impact of prior pregnancy history and AD-related genotype (PSEN2 knockout – KO) on late life cognitive function and neuropathology in aged female mice (10-15 months-old). Behavioral task performance was assessed in wild-type (WT) and PSEN2 KO mice, both nulliparous (NP) and multiparous (MP), before immunofluorescence was conducted for expression of various gliosis and neuroplasticity related proteins. We hypothesized that the AD-related risk gene, PSEN2, and parity history would differentially impact cognitive function and neuropathology in late life.

### Open Field (OFT) Behavioral Testing of Anxiety-Like Behavior

Ambulation, locomotion, and exploratory behavior were assessed using the Open Field Test (OFT), as previously published (Knox et al., 2023). An infrared-sensitive motion detection system (SuperFlex Open Field, Omnitech Electronics) was used to assess exploratory behavior and locomotion. Each mouse was placed into an OFT Plexiglas chamber (40 L x 40 W x 30H cm) for 10 mins, while their movements were monitored with the Fusion v6.5 (Omnitech) software. Locomotor chambers were cleaned with water between mice.

Recorded OFT parameters included total distance traveled (cm), vertical activity counts, and horizontal activity counts in both the center and the perimeter zones. OFT chamber center versus perimeter preference was quantified to measure anxiety-like behaviors (Seibenhener & Wooten, 2015).

### T Maze (TM) Testing of Working Memory

The TM spontaneous alternation test is a land-based task used to test hippocampal-mediated spatial working memory in mice (Deacon & Rawlins, 2006), as previously used with 6-month-old PSEN2 KO mice (Knox et al., 2023). TM was conducted 2 days after OFT. Each TM test consisted of 9 total sessions per mouse, with 3 sessions per day for 3 days. Mice were challenged to spontaneously alternate between the left and right goal arms (30 x 10 cm) of a T-shaped epoxy-coated wooden maze (start alley 30 x 10 cm). The spontaneous alternation rate was calculated as the ratio between alternating choices and total number of choices during all trials, with 60% considered successful TM performance, as previously established in 6-month-old female PSEN2 KO mice (Knox et al., 2023).

### Brain Tissue Collection and Processing

Mice were euthanized approximately 1 h after the last TM session, and brains were collected and hemi- sected longitudinally. The left-brain hemisphere was flash frozen and preserved for unrelated studies.

The right-brain hemisphere was fixed for 24 h in 4% paraformaldehyde at 4°C, then stored in 30% sucrose with 0.05% sodium azide for cryoprotection and preservation at 4°C for this study. The sucrose-submerged right-brain hemisphere was then flash frozen, before cryosectioning (Leica CM1850) into 30-μm thick slices from the prefrontal cortex (AP from Bregma: 1.98 to 1.70 mm) and the dorsal hippocampus (AP from Bregma: -1.58 to -2.10 mm). A total of 20 prefrontal cortex sections and 20 rostral and 20 caudal hippocampal sections were collected sequentially from each mouse, with 2 sequential sections per brain region/mouse mounted onto SuperFrost slides (Fisher). The slides were stored at -80°C until immunofluorescent processing.

### Immunofluorescence

Slides were washed with 1X PBS (3 x 5 min) before undergoing antigen retrieval with citrate buffer (#ab93678; Abcam) heated at 95 °C for 10 min, then resting at room temperature for 20 min. GAD65/67 and PV underwent antigen retrieval for 20 min at 95 °C, then 40 min at room temperature. The slides were then washed with dH_2_O (3 x 2 min). Antibodies for all molecular targets are detailed in Table 1.

Slides immunolabeled for GAD65/67 and PV were permeabilized with 0.3% Triton X-100 in 1X PBS for 30 min before undergoing blocking incubation with 10% goat serum with 0.03% Triton-X 100 in 1X PBS with 200 mL coverwells for 2 hr at RT. For other markers, a different blocking reagent (10% goat serum with 5% Triton-X 100 in 1X PBS) was applied for 2 hr at RT. Next, the GAD65/67 and PV antibodies were applied to the slides in a 1% goat serum, and 0.03% Triton-X in 1X PBS solution with coverwells. Other antibodies were applied with 5% Goat Serum in 1X PBS. Slides for GAD65/67, PV, BDNF, and Plp underwent primary antibody incubation at 4 °C overnight, whereas slides for NeuN, EAAT2, and Iba1 were incubated for 2 hr at RT. The next day, the slides were washed with 1X PBS for 3 x 5 min and incubated with either anti-rabbit secondary antibody or anti-mouse secondary antibody, with addition of conjugated GFAP. For GAD65/67 and PV, the secondary antibody was applied in 1% goat serum, 0.03% Triton-X in 1X PBS. For remaining markers, secondary incubation was conducted in 5% goat serum in 1X PBS. Slides for GAD65/67, PV, BDNF, Plp, and GFAP underwent secondary antibody incubation for 2 hr at RT, whereas slides for NeuN, EAAT2, and Iba1 were incubated at 4 °C overnight.

A final wash was done with 1X PBS (3 x 5 min) before the slides were coverslipped with Prolong Gold Antifade with DAPI (#P36934; Thermo Fisher).

The hippocampus, largely associated with the process of episodic memory, not only exhibits neuroplasticity with pregnancy (Pawluski & Galea, 2006), but also experiences significant neurodegeneration in AD. Moreover, the broader limbic system and prefrontal cortex play a critical role in higher brain function, mediating cognitive control, emotion, and memory. There appears to be an effect of pregnancy on electrophysiology and synaptic activity in the hippocampal CA1 region, the basolateral amygdala (BLA), and the medial prefrontal cortex (MPFC) (Anastasiades & Carter, 2021; Bin & Luo, 2025). Two photomicrographs were thus captured from each animal for these brain regions (prefrontal cortex, subregions of the dorsal hippocampus, the somatosensory cortex overlaying CA1 of the dorsal hippocampus, and the basolateral amygdala) with a fluorescent microscope (Leica DM-6) with a 20x objective (80x final magnification) a minimum of 24 hr after coverslip mounting. Acquisition settings were held constant throughout. Plp, NeuN, PV, GAD65/67, BDNF, EAAT2, Iba1, and GFAP expression was reported as percent total field area with fluorescent signal for the molecular marker of interest and quantified with ImageJ (Fiji).

### Statistical Analyses

Behavioral performance on the OFT and TM, as well as immunofluorescent protein expression, were compared by two-way ANOVA (parity x genotype), with post-hoc Dunnett’s test when applicable. All analyses were conducted with GraphPad Prism v10.0 with p < 0.05 considered significant.

## RESULTS

### Pregnancy history does not impact short-term spatial working memory, but does significantly increase performance task recall latency of aged female mice

Performance of 10-15-month-old WT versus PSEN2 KO mice was assessed on the T Maze (TM) continuous alternation task (Figure 2) because we have reported performance deficits on this task in 6-month-old PSEN2 KO female mice (Knox et al., 2023). The number of TM response errors did not significantly differ by genotype nor parity history (Figure 2A). All groups had a response rate similar to typical alternation rates of healthy WT mice, which is reported as an error rate equal to or less than 40% (Deacon & Rawlins, 2006). Similarly, there was no significant difference between the number of responders, defined as the mice that met this healthy WT criteria, between the various groups (Fig 2B).

**Figure 2.**
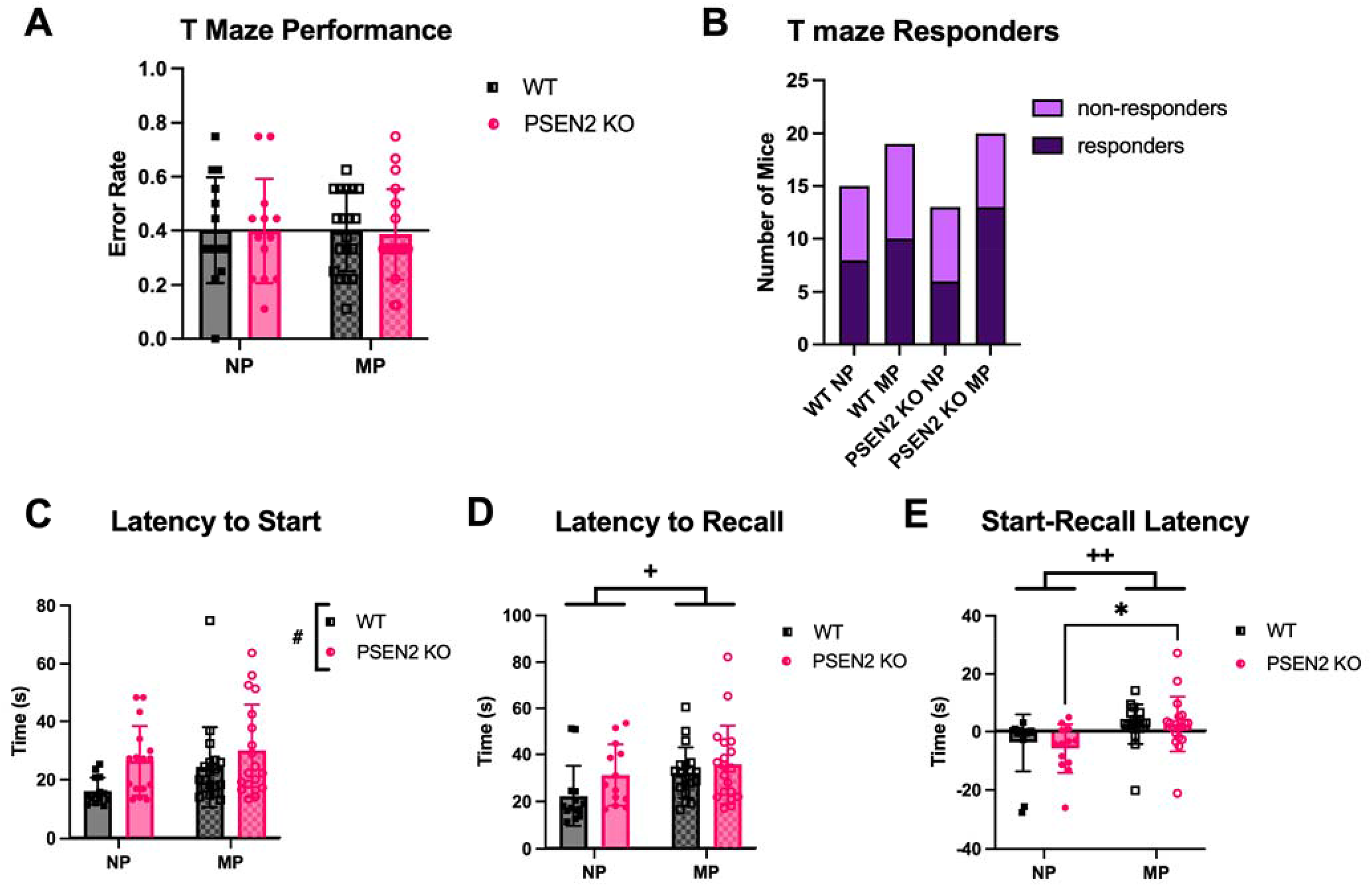
Nulliparous (NP) and multiparous (MP) female mice with loss of normal PSEN2 function were challenged in the spatial working memory T maze (TM) task to assess cognitive function at age 10-15 months-old. A) TM spontaneous alternation error rate was no different in NP and MP mice. All groups performed to the same degree as NP WT mice (defined as error rate <40%), indicated by the horizontal line. B) The numbers of mice in each experimental group that met responder criteria of >60% correct responses were assessed and neither genotype nor pregnancy history influenced responder rate. C) Genotype exerted a main effect on the latency to start, where PSEN2 KO mice exhibited a longer latency than WT mice. D) There was a main effect of parity on the latency to recall and the E) start- recall latency, where multiparous females took a significantly longer time to complete the task. The latency to start and latency to recall are defined as the time it took for each mouse to choose a goal arm, either on the initial or secondary trial respectively. The start-recall latency is the difference between these values. PSEN2 KO multiparous mice had a significantly higher start-recall latency compared to PSEN2 KO NP mice. This means the PSEN2 KO multiparous mice took longer to complete the second arm choice, relative to the initial trial, whereas the PSEN2 KO NP mice exhibited the opposite trend. All comparisons were analyzed with two-way ANOVA. # indicates main effect of genotype, + indicates main effect of parity, * indicates individual group differences, (*/+/# p<0.05, ++ p<0.01).

Genotype significantly influenced the latency to initiate the task (i.e., to “start” the trial; (F (1, 64) = 6.63, p=0.012), where PSEN2 KO mice took significantly more time to complete the initial goal arm choice, compared to the WT mice (Fig 2C). Parity significantly increased the recall latency (F(1, 61) = 4.44, p=0.04), meaning that multiparous females took significantly longer to initiate the secondary arm decisions and/or recall previous arm choice (Fig 2D). Since response accuracy was not affected in the TM, these increased latencies may indicate elevated stress or decreased motivation and/or interest in the task. Furthermore, parity also influenced the delay between trial start and decision (i.e. the start- recall latency, Fig 2E; F(1, 61) = 11.61, p=0.001). Post-hoc analysis revealed that PSEN2 KO multiparous females had a significantly greater start-recall latency than PSEN2 KO nulliparous females (p=0.034), meaning the nulliparous mice took longer to choose a goal arm in the second recall trial, relative to the initial start trial, compared to the genotype-matched multiparous mice. Overall, neither genotype nor parity affected hippocampal-mediated spatial working memory capacity as demonstrated by no difference in error rates between groups. However, parity history may be associated with memory and recall delays regardless of genotype.

### Pregnancy history and presenilin 2 deficiency exert differing long-term effects on open field locomotion and anxiety-like behaviors

Both exploratory and ambulatory behaviors were assessed in the OFT, revealing that both genotype and parity significantly influenced locomotor activity (Fig 3A). Specifically, AD-related genotype was associated with significantly decreased total distance traveled (Fig 3B; genotype: F(1, 61) = 41.6, p<0.0001) and total horizontal activity (Fig 3C; genotype: F(1, 61) = 27.2, p<0.0001) in the OFT.

**Figure 3.**
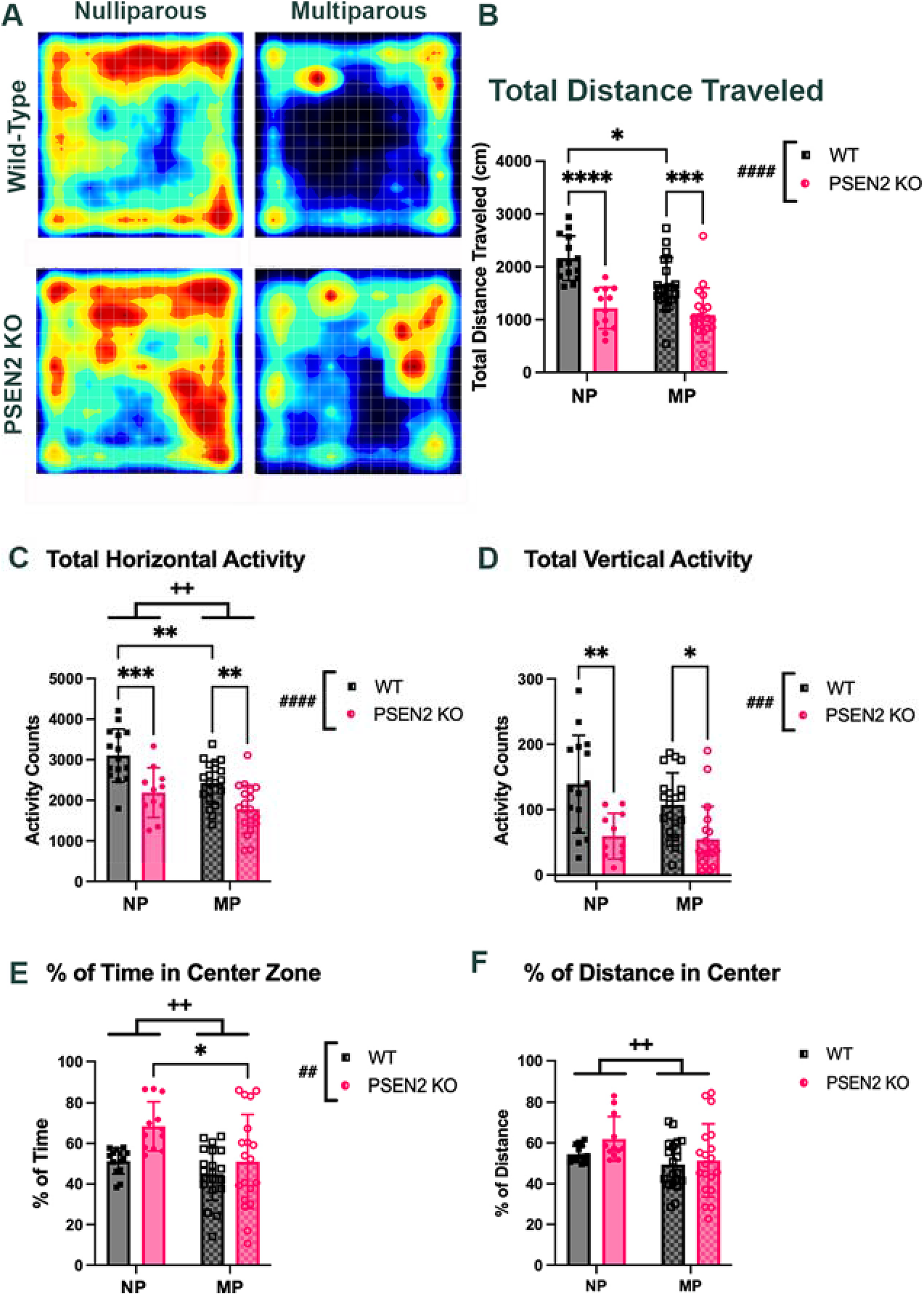
Genotype and parity status of 10-15-month-old WT and PSEN2 KO female mice exerted differing effects on Open Field Task parameters for ambulation, locomotion, and anxiety-like behaviors. PSEN2 KO and WT mice with (multiparous – MP) and without (nulliparous – NP) pregnancy history were challenged in a non-habituated open field test. A) Representative individual mouse heat map images of open field maze exploration over 10 minutes to represent the average behavioral activity and preference zones for each experimental condition. B) There was a significant effect of genotype and parity in total distance travelled, with PSEN2 KO mice exhibiting decreased distance travelled compared to WT, in both NP and MP groups. Furthermore, for the WT groups, MP mice had significantly decreased distance traveled compared to NP mice. Similar trends for total distance travelled was observed for C) total horizontal activity. D) For total vertical activity, PSEN2 KO genotype significantly influenced activity counts. For both NP and MP groups, PSEN2 KO mice exhibited decreased vertical activity compared to parous-matched WT mice. E), F) Analysis of center versus perimeter open field maze exploration, which serves as an indicator for thigmotaxis, was determined with percentage of time and distance in the center zone. E) With percentage of time in the center, both parity and genotype exerted a significant main effect with both multiparity and PSEN2 KO status reducing center preference. This effect was most evident in the PSEN2 KO NP versus the genotype-matched MP mice. F) For percentage of distance in the center, MP females traveled less distance in the center compared to NP, however there were no individual group differences. All comparisons were analyzed with two-way ANOVA. # indicates main effect of genotype, + indicates main effect of parity, * indicates individual group differences, (*/+ p<0.05, **/++/## p<0.01, ***/### p<0.001, ****/#### p<0.0001).

Further, pregnancy history was also associated with reduced OFT total distance traveled (Fig 3B; F(1, 61) = 6.68, p=0.012) and reduced horizontal activity (Fig 3C; F(1, 61) = 13.0, p=0.0006). Post-hoc analyses revealed that for both total distance traveled and total horizontal activity, WT mice significantly differed from PSEN2 KO in both nulliparous (total distance; p<0.0001, horizontal activity; p=0.005) and multiparous mice (total distance; p=0.0009, horizontal activity; p=0.044). WT nulliparous mice also traveled significantly more total distance and had more horizontal exploratory activity versus WT multiparous mice (total distance; p=0.018, horizontal activity; p=0.0009), which was not observed in PSEN2 KO mice. While genotype and parity exert effects on locomotion, such that multiparity and loss of normal PSEN2 function each reduced ambulation independently, there was no genotype x parity interaction in either total distance traveled or horizontal activity (Figs 3B and 3C). Additionally, loss of normal PSEN2 function was associated with generally reduced total vertical activity relative to parity history-matched WT mice (Fig 3D; F(1, 61) = 22.7, p<0.0001). Vertical activity indicates exploratory behaviors such as rearing, where the mouse stands on its hind paws, suggesting that the PSEN2 KO mice exhibit reduced motivation to explore novel environments. Post-hoc analyses revealed that the WT and PSEN2 KO females were significantly different in both the nulliparous (p=0.0019) and the multiparous groups (p=0.015; Fig 3D) reflecting a general effect in aged PSEN2 null aged females that is consistent with and extends our prior findings (Knox et al., 2023).

The percentage of total time or distance in the OFT center can indicate anxiety-like behaviors. There was a main effect of parity, where multiparous mice spent more time (Fig 3E; F(1, 61) = 10.1, p=0.002) and distance (Fig 3F; F(1, 61) = 7.13, p=0.001) in the chamber periphery, indicating a more anxious-like phenotype. Post-hoc analysis revealed that PSEN2 KO multiparous mice spent significantly less of their total time in the OFT center versus PSEN2 KO nulliparous mice (Fig 3E; p=0.018; Fig 3E). There was also a main effect of genotype on percentage of time in the center (Fig 3E; F(1, 61) = 7.37, p=0.009), with PSEN2 KO mice spending more of their total exploration time in the OFT center; however, there was no genotype x parity interaction. Overall, PSEN2 genotype and parity did not synergistically worsen OFT performance and instead seemed to moderate cognitive performance in differing ways; genotype more significantly influenced motor activity and exploratory behaviors (Fig 3B-3D), whereas parity had a larger influence on anxiety-like behavior (Figs 3E-3F).

### Myelin density and neuronal structure are differentially impacted by loss of normal PSEN2 function and pregnancy history

Proteolipid protein (Plp) is the primary protein component of the central nervous system (CNS) myelin sheath formed by oligodendrocytes (*PLP1 Proteolipid Protein 1 [Homo Sapiens (Human)] - Gene - NCBI*, n.d.). Importantly, Plp expression is elevated in aged female mouse hippocampus and Plp up- regulation in oligodendrocytes improves cognition in old female mice, compared to male counterparts (Gadek et al., 2025). Given that myelin damage and impairment has been clinically reported in AD (Papuć & Rejdak, 2018), we hypothesized that PSEN2 deficiency and pregnancy history would influence this protein expression in aged female mice (Fig 4). There was a main effect of genotype in the hippocampal dentate gyrus (DG) region (Fig 5B; F (1, 40) = 9.87, p=0.003), with PSEN2 KO mice exhibiting more Plp expression versus WT. Post-hoc analyses reported DG Plp levels of multiparous PSEN2 KO mice were significantly elevated compared to multiparous WT mice (p=0.04). PSEN2 KO mice showed greater PLP expression in CA3 versus WT mice (Fig 5C; F(1, 38) = 10.64, p=0.002). Plp expression was significantly impacted by parity in the hippocampal CA1 (Fig 4D; F(1, 40) = 5.29 p=0.027) and medial prefrontal cortex (Fig 4E; MPFC; F(1, 36) = 4.43, p=0.042), with multiparous females exhibiting decreased Plp. There was a parity x genotype interaction on Plp expression in MPFC (F(1,36) = 4.74, p=0.036), with significantly greater expression in nulliparous WT mice relative to multiparous WT mice (p= 0.007), but this parity effect disappeared in the PSEN2 KO groups. This suggests that parity history does not influence medial prefrontal cortex Plp expression in PSEN2 KO mice, a region heavily associated with cognitive control and learning. There was a significant genotype x parity interaction in somatosensory cortex (Fig 4F; F(1, 39) = 5.89, p=0.02), but no significant post-hoc differences between the assessed groups. Lastly, there were no changes in PLP expression in BLA because of parity (F(1, 36) = 0.67, p>0.05) or genotype (F(1, 36) = 1.61, p>0.05; Fig 5G).

**Figure 4.**
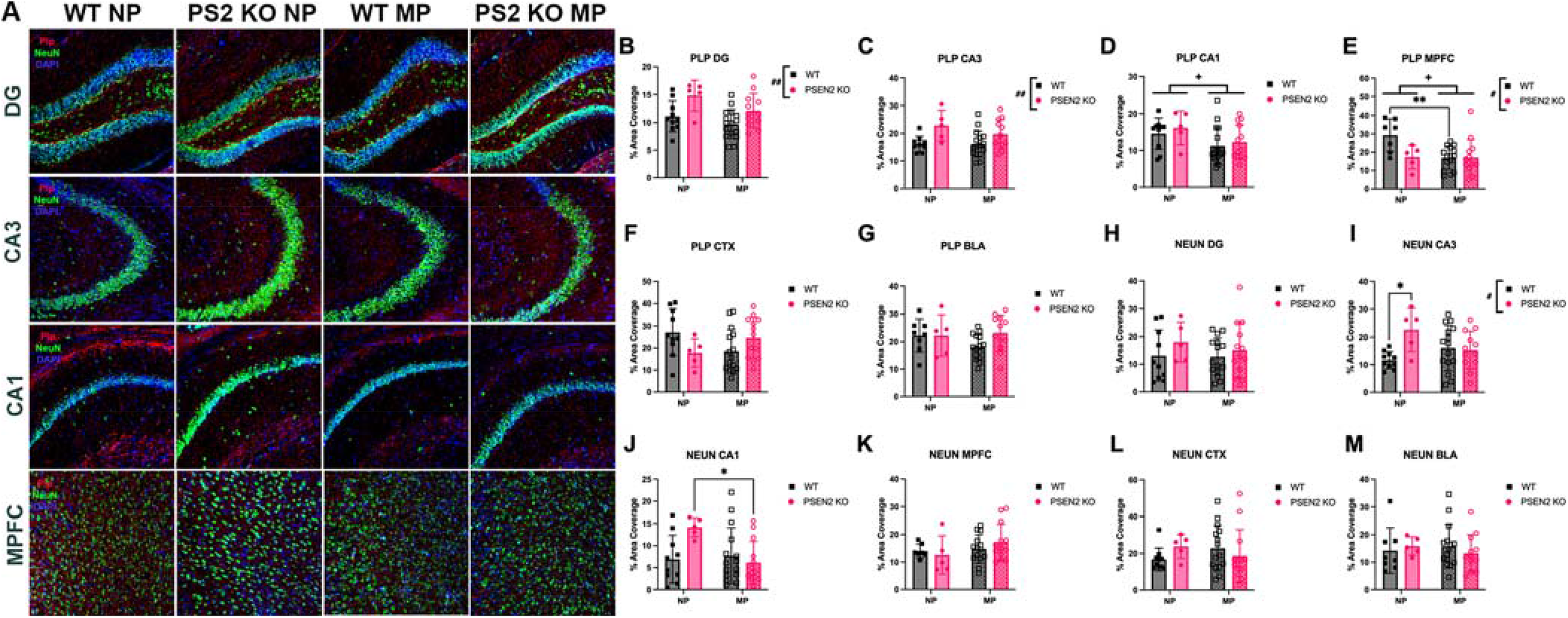
Proteolipid Protein (Plp) and Neuronal Nuclei (NeuN) expression in 10-15-month-old WT and PSEN2 KO mice with (multiparous - MP) and without (nulliparous - NP) pregnancy history after learning and memory behavioral testing. Myelin sheath integrity (as measured by Plp; red; B to G) and mature neuron density (as measured by NeuN; green; H) to M)) was analyzed in dorsal hippocampus (Dentate Gyrus (DG), CA3, CA1; B to D, H to J), medial prefrontal cortex (MPFC; E, K), somatosensory cortex (CTX; F, L), and basolateral amygdala (BLA; G, M). DAPI nuclear counterstain was also applied (blue). A) Representative photomicrographs of immunofluorescent labeling of Plp and NeuN in mouse brain in WT and PSEN2 KO NP and MP mice. PSEN2 KO genotype significantly influenced myelin sheath structure in the B) DG and C) CA3 region, with PSEN 2 KO females displaying increased Plp expression in the DG when compared to parous-matched WT. D) There were significant parity-induced decreases in Plp expression in the CA1 region, with MP females exhibiting lower Plp area coverage. E) In the MPFC, Plp expression was significantly changed by both parity and genotype, but there was no parity x genotype interaction. WT NP females had significantly greater Plp coverage in the MPFC versus WT MP females. There were no significant differences in Plp levels in the F) CTX and the G) BLA. For NeuN signaling in the I) hippocampal CA3 region, PSEN2 KO mice had increased NeuN expression compared to the WT mice, specifically in the NP group. J) there was a significant parity x genotype interaction in CA1 region, with the PSEN2 NP mice having greater neuronal density compared to the WT MP mice, but this trend was not observed in genotype or parity-matched pairs. The H) DG, K) MPFC, L) CTX, and M) BLA regions did not exhibit any significant differences in NeuN expression. All comparisons were analyzed with two-way ANOVA. # indicates main effect of genotype, + indicates main effect of parity, * indicates individual group differences, (*/+/# p<0.05, **/## p<0.01).

**Figure 5.**
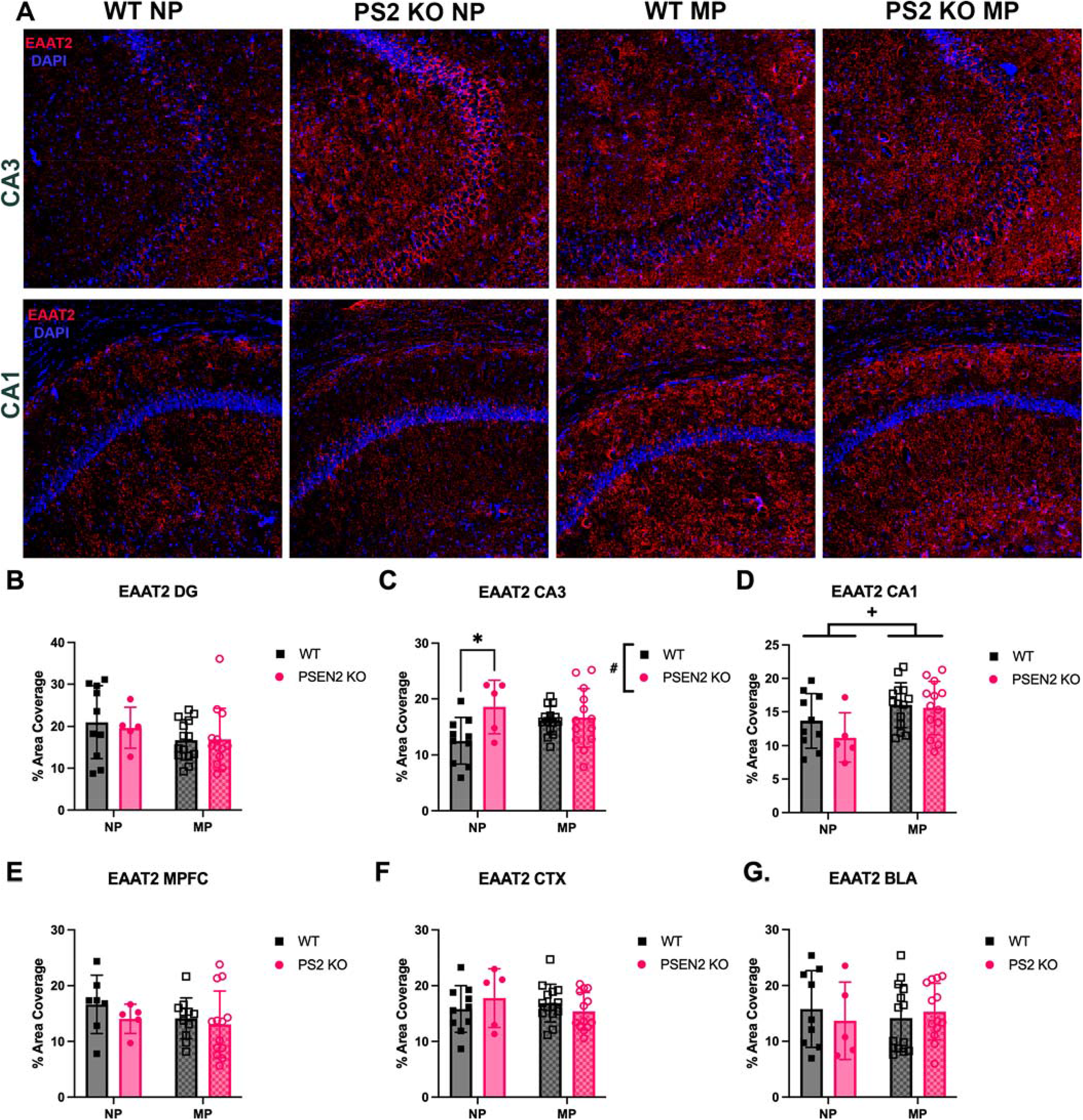
Excitatory amino acid transporter 2 (EAAT2) expression in 10-15-month-old WT and PSEN2 KO female mice with (multiparous - MP) and without (nulliparous - NP) pregnancy history after the conclusion of spatial memory testing. EAAT2 is responsible for glutamate reuptake from the synaptic cleft and serves to prevent excitotoxicity. EAAT2 expression (red) was analyzed in dorsal hippocampus (Dentate Gyrus (DG), CA3, CA1; B to D), medial prefrontal cortex (MPFC; E), somatosensory cortex (CTX; F), and basolateral amygdala (BLA; G). DAPI nuclear counterstain was applied (blue). A) Representative photomicrographs of immunofluorescent staining of EAAT2 in mouse brain in WT and PSEN2 KO NP and MP mice. B) There were no differences in EAAT2 staining in DG. C) EAAT2 was significantly influenced by genotype in the CA3 region. Post-hoc analysis revealed that female PSEN2 KO mice had greater EAAT2 expression versus WT in the NP groups specifically. D) In the CA1 region, MP females had significantly more EAAT2 expression compared to NP, regardless of genotype, indicating greater inhibitory tone with pregnancy history. The B) DG, E) MPFC, F) CTX, and G) BLA did not demonstrate any significant differences in EAAT2 signal because of parity or genotype status. All comparisons were analyzed with two-way ANOVA. # indicates main effect of genotype, + indicates main effect of parity, * indicates individual group differences, (*/+/# p<0.05).

We also assessed neuronal nuclei (NeuN) expression, which identifies post-mitotic, mature neurons in the CNS (Moon et al., 2026). There was differential NeuN expression in some, but not all, dorsal hippocampus regions (Fig 4H-4J). There was a genotype x parity interaction in both CA3 (Fig 4I; F(1, 38) = 6.48; p=0.015) and CA1 (Fig 4J; F(1, 39) = 6.05, p=0.018) such that the greater NeuN expression in the CA3 of nulliparous PSEN2 KO mice was not seen in the multiparous counterparts, suggesting pregnancy history may dampen PSEN2 KO associated changes in neuron density. Post-hoc analyses also demonstrated greater NeuN immunoreactivity in CA1 in the nulliparous versus multiparous PSEN 2 KO mice (p= 0.029), suggesting that pregnancy may differentially impact individual brain region neuronal density well into late life. All other regions (hippocampal DG, MPFC, cortex, and BLA; Fig 4H, 4K-M) demonstrated no differences in NeuN expression due to genotype or parity.

### Pregnancy history is associated with altered inhibitory/excitatory neurotransmitter protein expression in aged female mice that is not affected by PSEN2 KO genotype

We next sought to determine whether parity history could interact with an AD-related genotype to exert sustained modulation of the inhibitory/excitatory signaling balance in late adulthood. EAAT2, the excitatory amino acid transporter 2, clears glutamated from the synaptic cleft and is primarily expressed by astrocytes (Lin et al., 2012). EAAT2 expression was moderately affected by PSEN2 KO genotype and pregnancy history, mainly in the dorsal hippocampus. Genotype significantly impacted EAAT2 density in CA3 (Fig 5C; F(1, 38) = 5.55, p=0.024), with nulliparous PSEN2 KO mice having significantly greater EAAT2 immunoreactivity than nulliparous WT mice (p= 0.044), potentially indicating reduced excitatory transmission or astrocyte proliferation. The same trend was not seen in the multiparous group, suggesting that pregnancy history could blunt EAAT2 overexpression in PSEN2 KO. In CA1, EAAT2 expression was greatest in multiparous mice (Fig 5D; F(1, 38) = 7.32, p=0.01), but no genotype-related differences emerged. No other brain regions exhibited sustained effects of genotype or parity.

GABA, the major inhibitory neurotransmitter, is synthesized from glutamate in an enzymatic reaction catalyzed by glutamic acid decarboxylase (GAD), which exists in two isoforms with molecular weights of 65 and 67 kDa (GAD65/67) (Martin & Rimvall, 1993). Thus, GAD65/67 expression provides insight into inhibitory balance and GABA production in the brain. Multiparous females consistently had more GAD65/67 immunoreactivity in CA1 (Fig 6D; F(1, 40) = 6.15, p=0.017), MPFC (Fig 7E; F(1, 38) = 5.84, p=0.021), CTX (Fig 6F; F (1, 40) = 7.61, p=0.009), and BLA (Fig 5G; F(1, 38) = 4.77, p=0.035). In those same brain regions, post-hoc analyses revealed that the multiparous WT females had more GAD65/67 immunoreactivity compared to nulliparous WT females (CA1; p= 0.03, MPFC; p=0.035, CTX; p=0.01, BLA; p=0.046) but this post-hoc effect was absent in PSEN2 KO mice. The remaining hippocampal DG and CA3 regions (Fig 6B, C) did not demonstrate any significant differences in GAD65/67 expression across the groups. Overall, pregnancy history exerts a broad and consistent influence on inhibitory neurotransmission that is not further altered by AD-related genotype.

**Figure 6.**
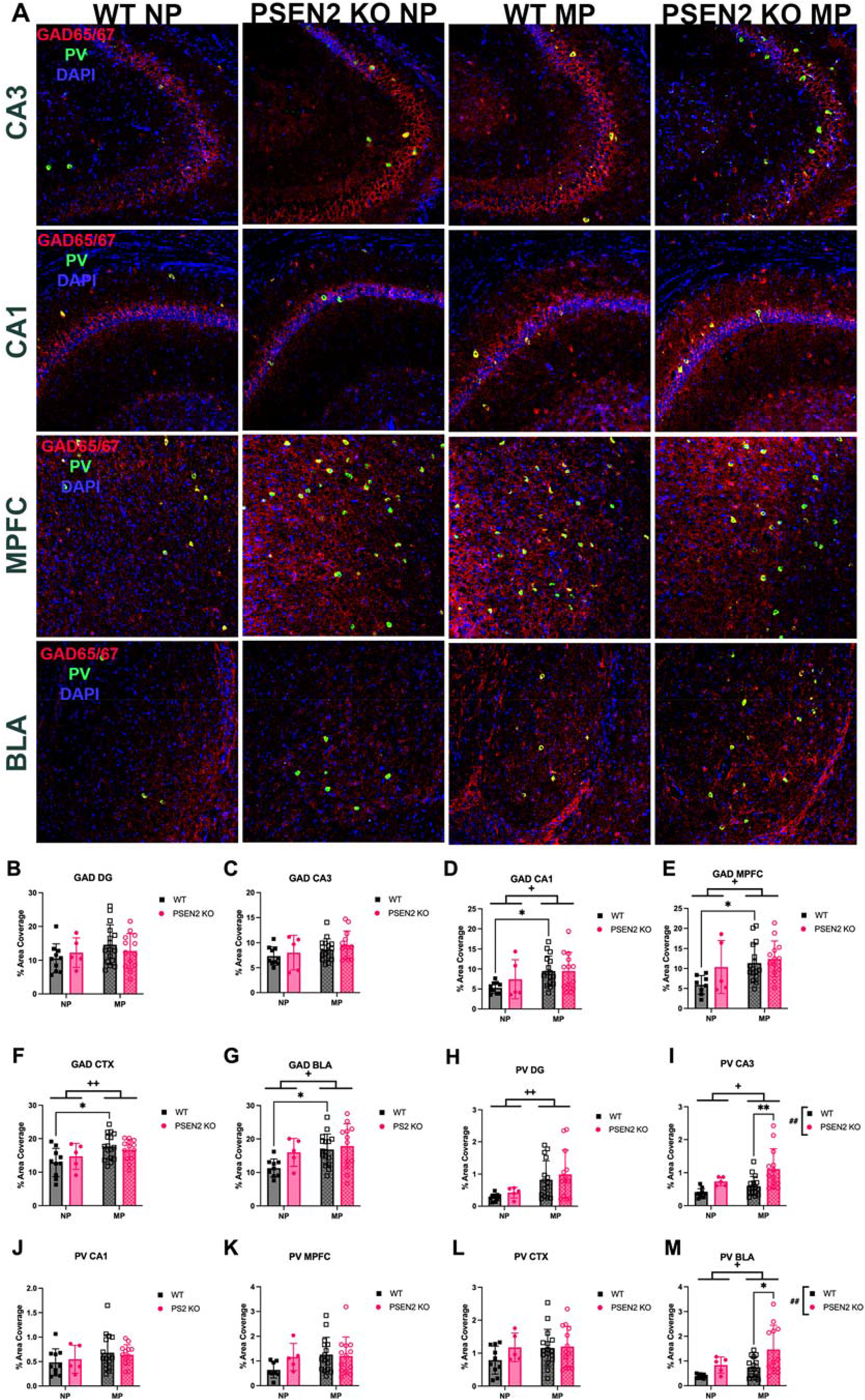
Inhibitory signaling measured via glutamic acid decarboxylase 65/67 (GAD65/67) and parvalbumin (PV) expression in 10-15-month-old WT and PSEN2 KO female mice with (multiparous - MP) and without (nulliparous - NP) pregnancy history after the conclusion of learning and memory behavioral testing. GAD65/67-mediated GABA synthesis (red; B to G) and inhibitory PV interneuron density (green; H) to M)) was analyzed in dorsal hippocampus (Dentate Gyrus (DG), CA3, CA1; B to D, H to J), medial prefrontal cortex (MPFC; E, K), somatosensory cortex (CTX; F, L), and basolateral amygdala (BLA; G, M). DAPI nuclear counterstain was also applied (blue). A) Representative photomicrographs of immunofluorescent labeling of GAD65/67 and PV-positive cells in the brains of WT and PSEN2 KO NP and MP mice. Parity influence on GAD65/67 expression throughout various brain regions in MP females in the D) CA1, E) MPFC, F) CTX, and G) BLA, compared to NP. Within these brain regions, specifically the WT MP females had consistently more GAD65/67 expression compared to genotype-matched NP mice. B) DG and C) CA3 did not experience any parity or genotype-mediated differences in GAD65/67 expression. Looking at PV interneuron coverage, the H) DG exhibited a main effect of pregnancy, with MP females displaying greater PV signal compared to NP, although there were no individual group differences. I) CA3 and M) BLA exhibited similar trends; both parity and genotype exerted a significant main effect, with PSEN2 KO females showing increased PV expression compared to WT, specifically in the MP groups. Remaining brain regions (J - L) had no difference in PV expression across groups. All comparisons were analyzed with two-way ANOVA. # indicates main effect of genotype, + indicates main effect of parity, * indicates individual group differences, (*/+ p<0.05, **/++/## p<0.01).

**Figure 7.**
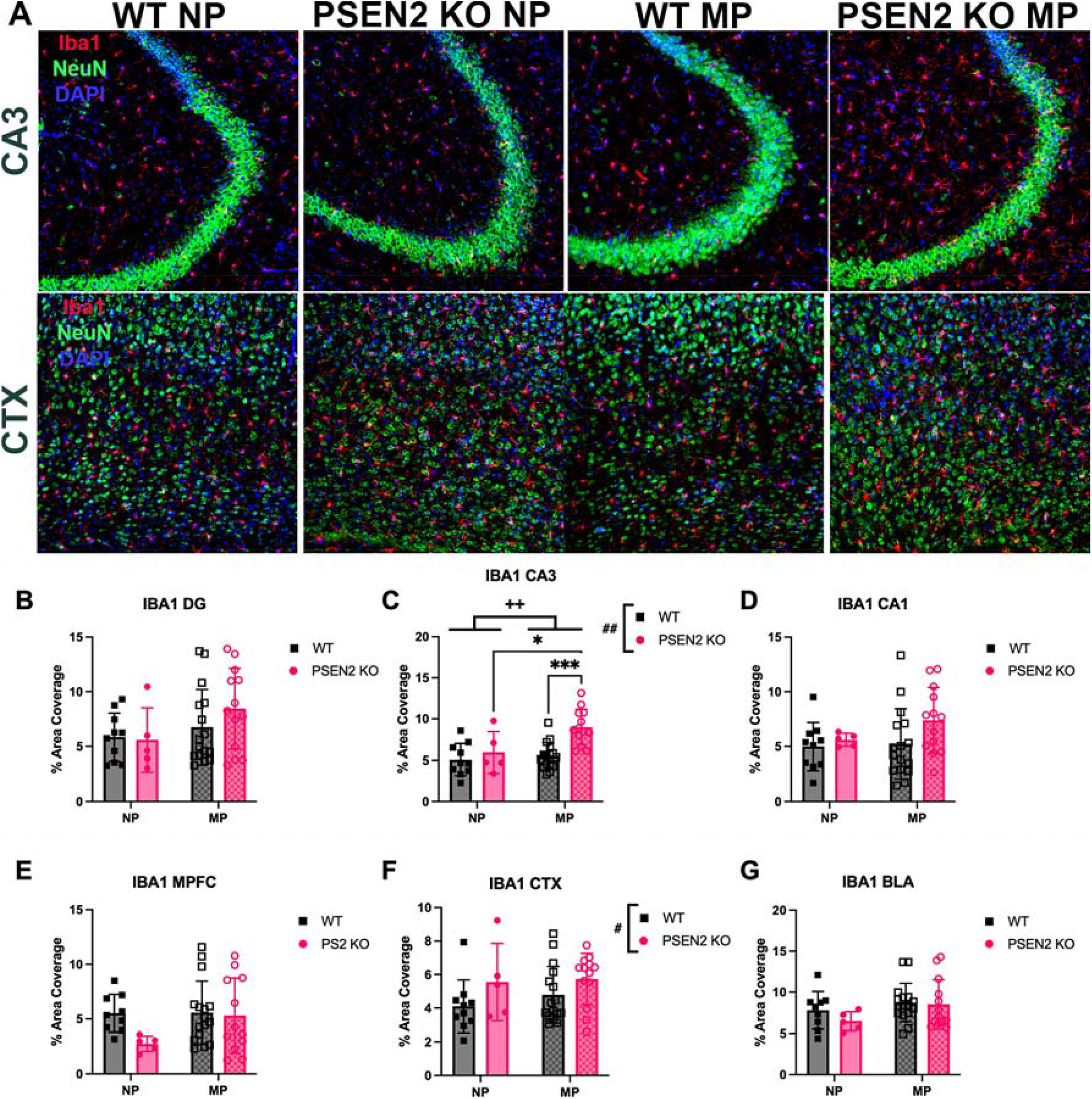
Aged (10-15-month-old) PSEN2 KO multiparous female mice exhibited greater reactive microgliosis in area CA3 of dorsal hippocampus relative to both PSEN2 nulliparous (NP) and WT multiparous (MP) animals. The expression of ionized calcium-binding adaptor protein (Iba1; red) was analyzed by immunofluorescence in the dorsal hippocampus (Dentate Gyrus (DG), CA3, CA1; B to D), medial prefrontal cortex (MPFC; E), somatosensory cortex (CTX; F) and basolateral amygdala (BLA; G) to mark microgliosis. NeuN (green) co-labeling and DAPI nuclear counterstain was also applied (blue). A) Representative photomicrographs of immunofluorescent labeling of Iba1 and NeuN in WT and PSEN2 KO NP and MP mice brains. In the C) hippocampal CA3 region, both parity history and PSEN2 genotype increased Iba1 signal, indicating elevated microglial activation. The microgliosis in the PSEN2 KO MP mice was significantly greater than both the WT multiparous mice and the PSEN2 KO NP mice. There was also a main effect of genotype in the F) cortex, with PSEN2 KO mice exhibiting elevated Iba1 expression compared to wildtype in both MP and NP groups. The remaining brain regions (B, D, E, G) displayed no differences in Iba1-associated microglia activation. All comparisons were analyzed with two-way ANOVA. # indicates main effect of genotype, + indicates main effect of parity, * indicates individual group differences, (*/# p<0.05, ++/## p<0.01, *** p<0.001).

Parvalbumin (PV) is a calcium binding protein expressed in the largest subpopulation of GABAergic inhibitory interneurons (Bjerke et al., 2020). PV provides an additional marker of the inhibitory signaling in discrete brain regions involved in cognitive function and anxiety-like behaviors. Parity history in aged female mice was associated with more PV expression in the hippocampal DG region (Fig 6H; F(1, 40) = 9.43, p=0.004). However, there were no post-hoc differences. Similarly, both parity and genotype was associated with greater immunoreactivity in CA3 (Fig 6I; parity: F(1, 40) = 5.43; p=0.025, genotype; F(1, 40) = 11.74, p=0.001) and BLA (Fig 6M; parity: F(1, 38) = 6.09, p=0.018, genotype: F(1, 38) = 8.42, p=0.006). There was no parity x genotype interaction in these regions. Specifically in the multiparous groups, PSEN2 KO mice had significantly more PV expression compared to parous-matched WT mice (CA3; p=0.004, BLA; p=0.012). These findings further suggest that pregnancy history can upregulate inhibitory signaling well into late life.

### Pregnancy history and/or loss of PSEN2 function changes long-term glial reactivity

Activated microglia upregulate the expression of ionized calcium-binding adapter molecule 1 (Iba1); its function assists with membrane ruffling and phagocytosis by microglia (Ohsawa et al., 2004). Thus, Iba1 serves as a marker for reactive microgliosis and neuroinflammation, albeit it cannot differentiate resident microglia from infiltrating macrophages. Nonetheless, we sought to examine whether prior pregnancy or AD-related loss of normal PSEN2 function would exert any lasting changes on Iba1 immunoreactivity in late life. Iba1 expression in hippocampal CA3 was higher with pregnancy history (Figure 7C; F(1, 40) = 7.32, p=0.01) and PSEN2 KO genotype (F(1, 40) = 11.20, p =0.002). Post-hoc analyses revealed that the PSEN2 KO multiparous females had significantly more Iba1 signal compared to both the WT multiparous females (p=0.0001) and the PSEN2 KO nulliparous females (p= 0.02). Similarly, there was also a main effect of genotype in the somatosensory cortex overlaying dorsal hippocampus (Fig 7F; F(1, 40) = 4.53, p=0.04), with PSEN2 KO females generally displaying increased microgliosis, however there were no pregnancy history-related differences. No other brain regions exhibited any significant differences in Iba1 microglial activation. This association between greater microgliosis and loss of normal PSEN2 function in late life aligns with previous literature demonstrating that loss of PSEN2 function may promote exaggerated inflammatory responses (Jayadev et al., 2010, 2013; Luquez et al., 2026).

Glial fibrillary acidic protein (GFAP) is the hallmark intermediate filament protein in astrocytes, a cell type critical for metabolic balance, energy homeostasis, and neuronal signaling in the CNS (Hol & Pekny, 2015). Reactive astrogliosis, a characteristic change in the function and morphology of astrocytes seen in neurological diseases such as AD, is associated with GFAP upregulation. We thus investigated whether pregnancy history or loss of PSEN2 function would alter astrogliosis. In all three dorsal hippocampus regions, pregnancy history significantly influenced GFAP expression. Multiparous females exhibited increased GFAP immunoreactivity, reflective of astrogliosis, in DG (Fig 8B; F(1, 40) = 15.61, p=0.0003), CA3 (Fig 8C; F(1, 40) = 20.0, p<0.0001), and CA1 (Fig 8D; F(1, 41)=6.903, p=0.012), compared to the nulliparous groups. In the DG, the WT multiparous group had significantly elevated GFAP signal compared to the WT nulliparous group (p=0.006), a trend that was preserved in CA3 (p=0.018) and CA1 (p=0.042). Furthermore, GFAP expression in the PSEN2 KO multiparous females was also significantly greater than that of the PSEN2 KO nulliparous females in the CA3 (p= 0.008). Thus, increased astrogliosis was broadly observed throughout the hippocampus due to pregnancy history. Other cortical and amygdala brain regions did not display visible GFAP immunoreactivity and were thus not analyzed.

**Figure 8.**
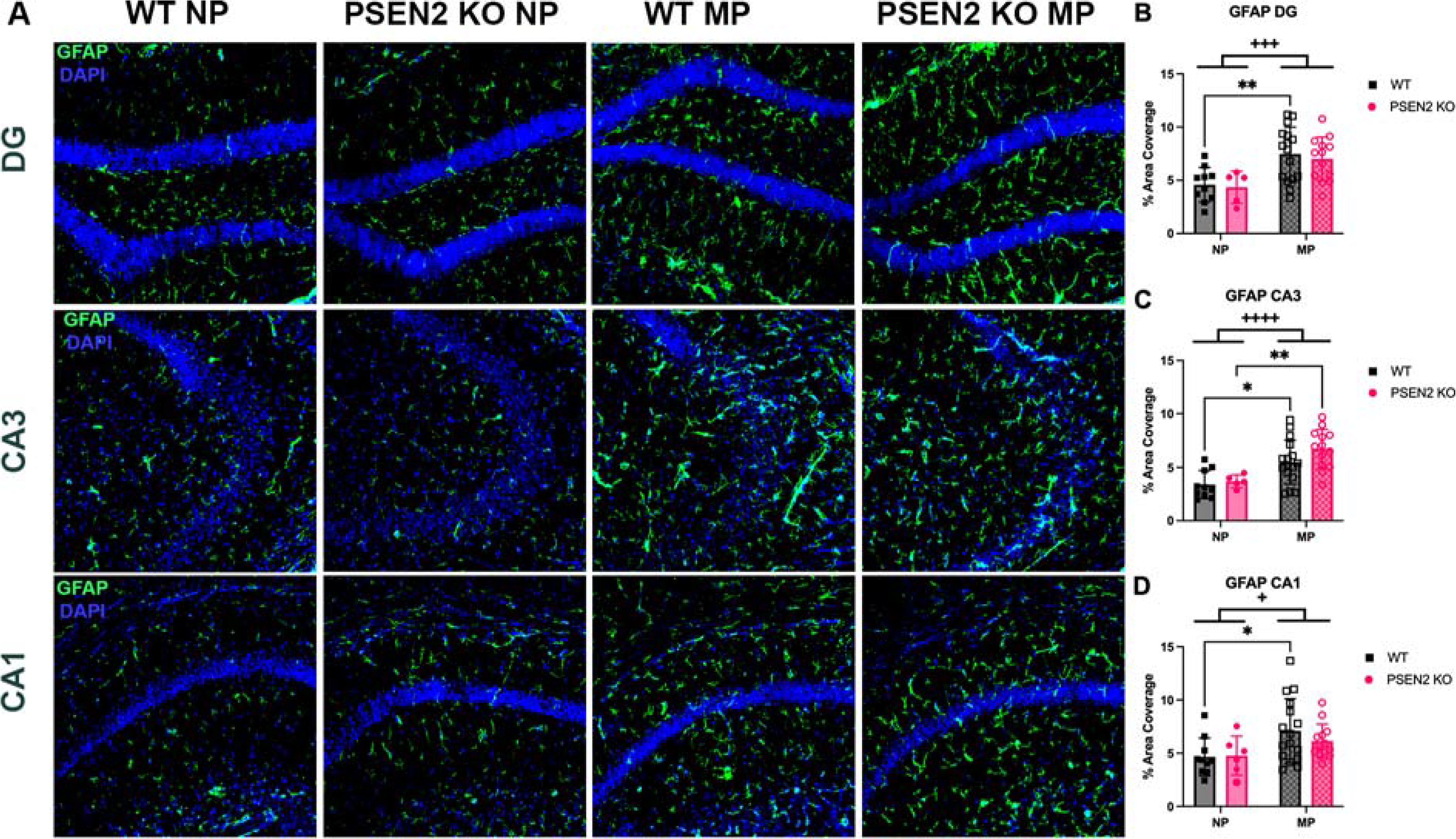
Aged (10-15-month-old) multiparous (MP) females demonstrated increased astrogliosis throughout various regions of the dorsal hippocampus compared to nulliparous (NP) females, irrespective of genotype. Glial fibrillary acidic protein (GFAP; green) expression was quantified via immunofluorescence to assess reactive astrocyte activation in the dorsal hippocampus (Dentate Gyrus (DG), CA3, CA1; B to D). DAPI nuclear counterstain was also applied (blue). A) Representative photomicrographs of immunofluorescent labeling of GFAP in WT and PSEN2 KO NP and MP mice hippocampal tissue. B) In the hippocampal DG region, pregnancy history exerted a significant effect on GFAP expression, with increased astrogliosis in MP WT versus NP WT females. A similar effect was seen in the C) CA3, with elevated GFAP levels observed in the MP females. In the CA3, both the WT and PSEN2 KO MP groups displayed significantly more GFAP immunofluorescence compared to genotype-matched NP animals. D) The same trend persisted in the CA1 region. For the WT animals, there was increased GFAP in the MP CA1 compared to NP. Remaining brain regions (CTX, MPFC, BLA) were not assessed for GFAP due to lack of signal across all animals. All comparisons were analyzed with two-way ANOVA. # indicates main effect of genotype, + indicates main effect of parity, * indicates individual group differences, (*/+ p<0.05, ** p<0.01, +++ p<0.001, ++++ p<0.0001).

### Pregnancy history and PSEN2 deficiency exert region-dependent differences in brain-derived neurotrophic factor (BDNF) expression in aged female hippocampus

BDNF plays an important role in neuronal survival and growth, modulating neurotransmitters, and influencing neuronal plasticity, which is crucial for memory and learning (Bathina & Das, 2015). Furthermore, decreased BDNF expression has been detected in AD, as well as other neurological conditions (Bathina & Das, 2015), and we wanted to investigate whether PSEN2 deficiency contributed to this reduction in BDNF. Moreover, we have previously demonstrated that repeated seizures, a means to consistently evoke chronic stress, leads to sex-specific decreases in BDNF expression in the hippocampal DG of 6-month-old female WT, but not PSEN2 KO, mice (Knox et al., 2023). In this study, a significant main effect of genotype revealed aged PSEN2 KO females exhibiting more BDNF signal in DG versus WT (Fig 9B; F(1, 40) = 5.03, p=0.031), regardless of parity status. This contradicts clinical findings and our prior work in younger animals; however, it is possible that elevated neurotrophic factors may reflect a compensatory response against early neuropathological remodeling associated with advanced age. In the CA3 region, parity exerted a significant main effect on BDNF levels (Fig 9C; F(1, 40) = 5.16, p=0.029). The multiparous group displayed increased BDNF compared to the nulliparous, specifically in WT multiparous versus nulliparous mice (p= 0.014). Thus, PSEN2 KO genotype and multiparity increased BDNF expression, albeit in a region-specific manner in the hippocampus. No other brain regions exhibited any significant BDNF expression differences.

**Figure 9.**
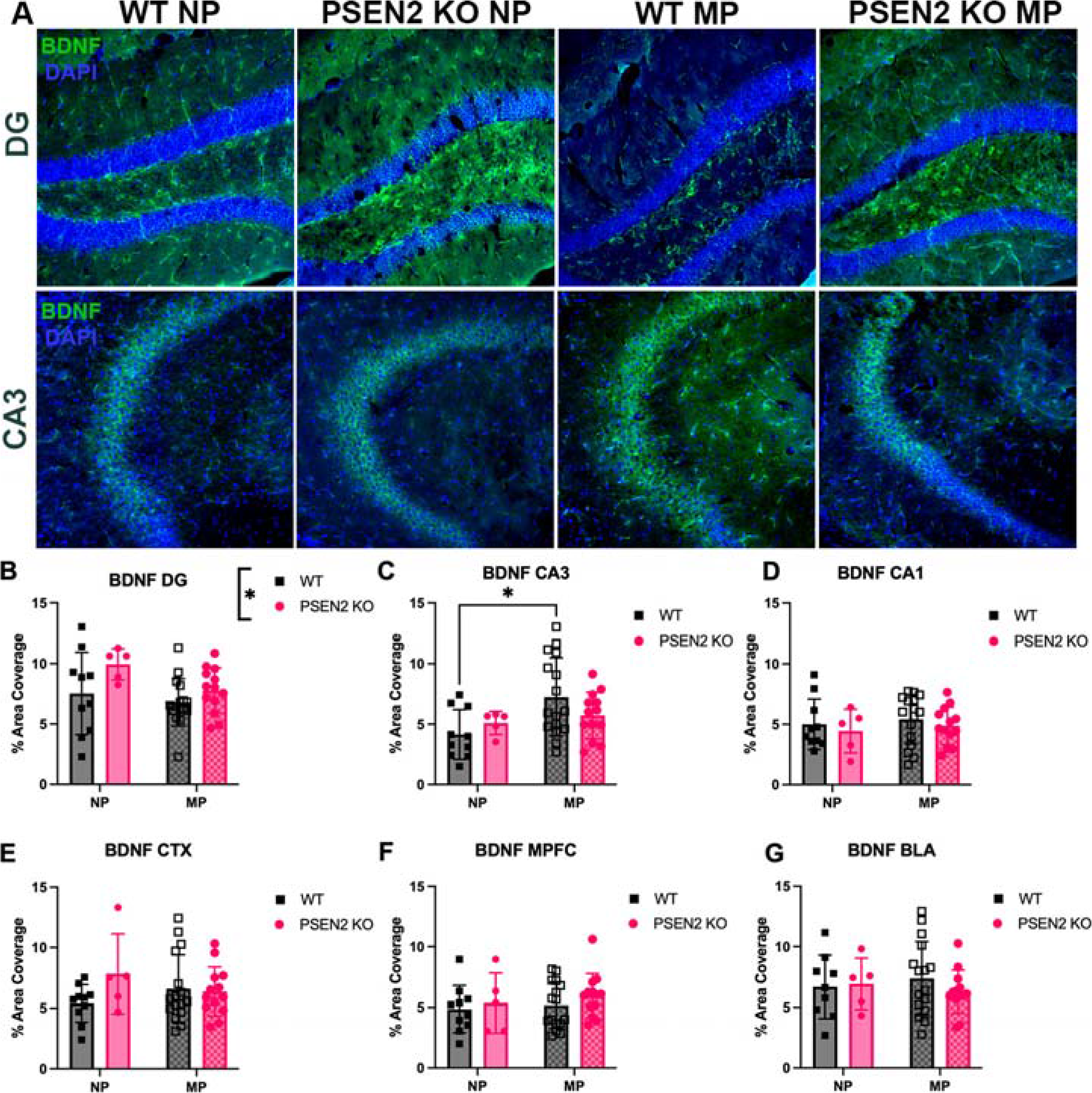
Brain-derived neurotrophic factor (BDNF) expression was increased in the DG of PSEN2 KO females, compared to WT, and the CA3 of multiparous (MP) females, compared to nulliparous (NP), indicating greater neuroplasticity in those groups. BDNF expression (red) was utilized to assess neuronal growth and survival via immunofluorescence in the dorsal hippocampus (Dentate Gyrus (DG), CA3, CA1; B to D), medial prefrontal cortex (MPFC; E), somatosensory cortex (CTX; F) and basolateral amygdala (BLA; G). DAPI nuclear counterstain was also applied (blue). A) Representative photomicrographs of immunofluorescent labeling of BDNF in WT and PSEN2 KO NP and MP mouse brain tissue. B) In the hippocampal DG region, genotype exerted a significant effect on BDNF expression, with increased levels in the PSEN2 groups versus WT females, regardless of parity status. C) In the CA3 region, there was a significant main effect of parity, with MP females exhibiting greater BDNF expression and increased neuronal growth. Specifically, BDNF expression was greater in the WT MP group compared to the WT NP group. The remaining brain regions (C, D, E, G) displayed no differences in BDNF expression. All comparisons were analyzed with two-way ANOVA. # indicates main effect of genotype, + indicates main effect of parity, * indicates individual group differences, (*/#/+ p<0.05).

## DISCUSSION

Females generally live longer but face more detrimental functional consequences of AD diagnosis. Defining how AD-associated risk factors interact with female-specific events is thus critical to improving healthy aging in women. In this study, while neither pregnancy history nor PSEN2 deficiency markedly impacted TM performance, the delayed latency to start the trial and recall previous information, and increased transition between behavioral tasks (start-recall latency) suggest generally reduced motivation. Both parity history and PSEN2 deficiency independently contributed to reduced OFT exploration. However, PSEN2 KO mice were generally less inhibited than WTs, spending more total time in the OFT center, albeit they were largely inactive. Pregnancy history increased thigmotaxis (reduced OFT center zone time), reflective of more anxiety-like behaviors. Although there was no impact to spatial working memory, the PSEN2 KO genotype alone consistently altered neuropsychiatric symptoms, namely anxiety and motivation, suggesting that PSEN2 may more meaningfully influence non-cognitive symptoms of AD (Shea et al., 2016; Yan et al., 2013). Further, we herein demonstrate that markers of neuroplasticity and neuroinflammation were meaningfully changed by pregnancy history and PSEN2 deficiency. Plp immunoreactivity was significantly impacted by both pregnancy history and PSEN2 deficiency across several cortical regions (Figure 3E and 3F). However, PSEN2 KO mice do not show late life amyloid-β plaque accumulation (Robinson-Cooper et al., 2026), suggesting that amyloid accumulation may not be the sole factor driving myelin dysfunction in AD (Kapasi et al., 2017; Moscoso et al., 2022). Pregnancy history was generally associated with decreased neuronal density, upregulation of inhibitory marker expression, and increased astrogliosis, which may contribute to the poor cognitive performance observed in the multiparous mice. Our study demonstrates that while pregnancy history does impart lasting changes in brain circuitry and cognitive processing with aging, it does not appear to synergistically interact with PSEN2 deficiency.

This present study is the first, to our knowledge, to evaluate the interaction between parity history and PSEN2 deficiency on late life cognitive performance and neuroplasticity-associated protein expression in the aged female brain. Cui and colleagues (2014) found that in APP23 mice, an amyloidogenic model, 12-month multiparous females displayed reduced Y-Maze alternation whereas in age-matched the WT groups, multiparous females experienced increased alternation compared to nulliparous. Another study found that primiparous (one birth) 12–13-month rodents with the hAPOEe4 genotype, a late onset AD-associated risk factor, were more likely to utilize a non-spatial strategy during the radial arm maze task, indicating decreased hippocampal recruitment to complete this task (Lee et al., 2024). This suggests that parity-mediated impairment to hippocampal-mediated spatial working memory performance may be mechanism-dependent and influenced by amyloidogenic (APP23) or lipid homeostasis (APOEe4) processes. Our study fills an important gap to demonstrate that loss of PSEN2 function and pregnancy history each seem to exert their own individual impacts on locomotor activities and anxiety-like behaviors.

Pregnancy history broadly altered inhibitory/excitatory signaling dynamics, measured by increased EAAT2, GAD65/67, and inhibitory PV interneuron protein expression. Pregnancy history was specifically associated with more GAD65/67 immunoreactivity in CA1, BLA, and cortical (somatosensory cortex and MPFC) regions of WT, but not PSEN2 KO female mice, suggesting that earlier reproductive events led to lasting remodeling of neuronal inhibition in a genotype-related manner. However, there was only a measurable shift in PV neuron expression in CA3 and BLA of multiparous PSEN2 KO mice, indicating that this inhibitory interneuron population was not responsible for the observed shift in GAD65/67. PV only identifies a subset of interneurons; thus upregulation of another GABAergic neuron subtype or increased inhibitory output from existing interneurons may have contributed. Elevated GAD expression in the lateral septum, accessory olfactory tract bed nucleus, and MPFC, brain regions heavily associated with anxiety and maternal behaviors, has been reported in postpartum mice compared to virgin females. (Ragan et al., 2022; Rodrı guez et al., 2004; Zhao et al., 2012). Furthermore, inhibitory cell upregulation is linked to anxiety-related behaviors, with chronic stress increasing prefrontal PV cell activity and chronic activation of these prefrontal PV cells inducing an anxiety-like phenotype in female mice (Page et al., 2019). This aligns closely with our findings, indicating that GABA modifications long after pregnancy are complex and pregnancy history may confer a sustained neurobiological effect on par with chronic stress, inducing inhibitory modulation to influence anxiety-like behaviors. Inhibitory signaling also displays vast regional differences in circuitry involvement and regulation. PV neurons are implicated in parahippocampal pathways facilitating episodic memory, the prefrontal cortex depends heavily on the inhibitory/excitatory balance for executive functioning, and GAD expression displays region-specific regulation in response to stress (Bjerke et al., 2020; Bowers et al., 1998; Ferguson & Gao, 2018). Thus, our presently observed regional differences in PV and GAD65/67 expression may reflect the diverse inhibitory neurotransmission contributions to behavioral and cognitive processes.

Pregnancy is a highly complex immunological event, with the maternal immune system undergoing specialized alterations to facilitate fetal development. We thus compared microglial and astroglial responses in aged females to investigate whether those changes persist into late life. PSEN2 KO mice showed increased Iba1 expression in CA3 and somatosensory cortex, supporting the conclusion that as the predominant γ-secretase in microglia (Jayadev et al., 2010), loss of normal PSEN2 function changes microglial reactivity well into late-life. Loss of normal PSEN2 function may disrupt microglia-mediated synaptic pruning in CA3, which could underlie the observed recall deficits in the TM (Fig 2E). Microglia display regional heterogeneity in density and morphology and are highly enriched in the hippocampus (Tan et al., 2020). Thus, variability in microglial distribution and physiology across different brain regions may have contributed to our presently observed Iba1 immunoreactivity differences. Microglia also interact with astrocytes, which regulate synaptic transmission and plasticity facilitating memory formation and anxiety-related behaviors (Cho et al., 2022; Wang et al., 2022). Parity history was herein associated with increased reactive astrogliosis across all three hippocampal brain regions, regardless of genotype. Altogether our study demonstrates sustained changes in hippocampal glial protein expression, with PSEN2 deficiency most dramatically changing Iba1 levels in CA3 and pregnancy driving sustained changes in GFAP.

Female cognitive resilience is increasingly documented in normal brain aging, with women generally experiencing reduced age-related declines in cognitive function and brain atrophy versus age-matched men (Ravndal et al., 2025; Xu et al., 2000). However, how pregnancy history uniquely influences female cognitive function in late life is less clear and subject to diverse socioeconomic confounds (M. M. Fox et al., 2026). Rodent past reproductive experience can influence hippocampal dendrite and neuron proliferation (Pawluski & Galea, 2006; Trainer et al., 2006), as well as increased synaptic plasticity (Duarte-Guterman et al., 2023) and enhanced learning and memory task performance (Kinsley et al., 1999; Zimberknopf et al., 2011; Macbeth et al., 2008). Herein, we found that pregnancy history did not adversely alter spatial working memory but was associated with changes in anxiety-like behaviors and locomotion. To our knowledge, our study had the advantage over prior studies that used younger rodents (oldest reported studies used mice 7 months-old) and tested animals much sooner after weaning (3 weeks; (Kinsley et al., 1999; Macbeth et al., 2008; Pawluski & Galea, 2006; Trainer et al., 2006)). Our study suggests that advanced age and extended delay from parturition (>7 weeks in our study) may modulate the initially neuroprotective effects of pregnancy. Thus, our present study carries substantial translational value to address the long-term effects of reproductive history well into late adulthood after reproductive senescence.

Female cognitive advantage may involve the X chromosome because female cells contain two X chromosomes, one of which is randomly active (Bourzac, 2025). An X chromosome genetic meta-analysis identified a X chromosome gene locus involved with secretory and endosomal function, and potentially downstream amyloid accumulation (Belloy et al., 2024). Given that PSEN2 KO mice lack amyloid pathology (Robinson-Cooper et al., 2026), this may explain the minimal impact of PSEN2 KO on cognitive outcomes. Furthermore, epigenetic gene silencing in the maternal X chromosome is associated with cognitive aging (Abdulai-Saiku et al., 2025). It is possible that the epigenetic modifications of pregnancy, in response to hormonal signaling, may silence genes associated with brain health (Andrawus et al., 2022). Plp1 undergoes age-induced re-activation of the silent X chromosome, and its expression in the hippocampus of aged females has been reported to positively correlate with cognitive function (Gadek et al., 2025). That parity history presently decreased hippocampal and medial prefrontal cortex Plp expression in conjunction with measurable deficits in behavioral performance on two tasks (i.e. increased latency to initiate goal choice in the TM (Fig 2D-E) and elevated anxiety-like behaviors in the OFT (Fig 3) aligns with this hypothesized role for Plp on cognitive function. Altogether, our study further points towards Plp, and X-chromosome associated markers as contributors to resilience in the aged female brain.

While our present study carries several impactful findings, we also acknowledge some limitations. First, most human studies report differential risk for AD and dementia outcomes based on the number of children, with more pregnancies associated with greater AD risk (Colucci et al., 2006; M. M. Fox et al., 2026; Jang et al., 2018). While socioeconomic factors could equally drive this difference, further experiments could benefit from the inclusion of primiparous mice (1 pregnancy) versus multiparous mice (multiple pregnancies) to more definitively establish whether reproductive history additively influences cognitive outcomes with aging. Another limitation is that the multiparous mice utilized were retired breeders, meaning that the exact number of pregnancies per female varied across individuals. Interindividual variability in total pregnancies could impact behavioral task performance and biomarker expression, especially if the positive correlation between the number of pregnancies and magnitude of AD risk holds true in animal models. The retired breeders consumed a high calorie chow to compensate for the energetically demanding requirements of frequent pregnancies and lactation, whereas the nulliparous mice did not. While it is likely that the caloric intake between the two groups closely met biological demands (Ladyman et al., 2018), we cannot rule out that subtle dietary factors could have also contributed to the presently observed neurological changes. Lastly, the hormone changes needed to sustain pregnancy are unparalleled by any other neuroendocrine events in a healthy woman’s life (Deems & Leuner, 2020). Future studies should prospectively monitor estrogen and progesterone levels to investigate whether short-term changes to sex hormone levels correlate to later cognitive impacts. Inclusion of sex hormones may more comprehensively define the differences that biological sex may induce on late-life cognitive function or AD risk. The effect of parity in the brain is both complex and diverse, further emphasizing the importance of studying female-specific experiences on late-life AD-related outcomes.

## Acknowledgements

The authors gratefully acknowledge the technical support of Stephanie Davidson and Kevin M. Knox. This work was supported by the Plein Center for Geriatric Pharmacy at the University of Washington (to JIF), K22AG082712 (to KEP) and R01AG067788 (to MBH). None of the authors have any conflicts of interest.

---

**Supplemental Table 1.**
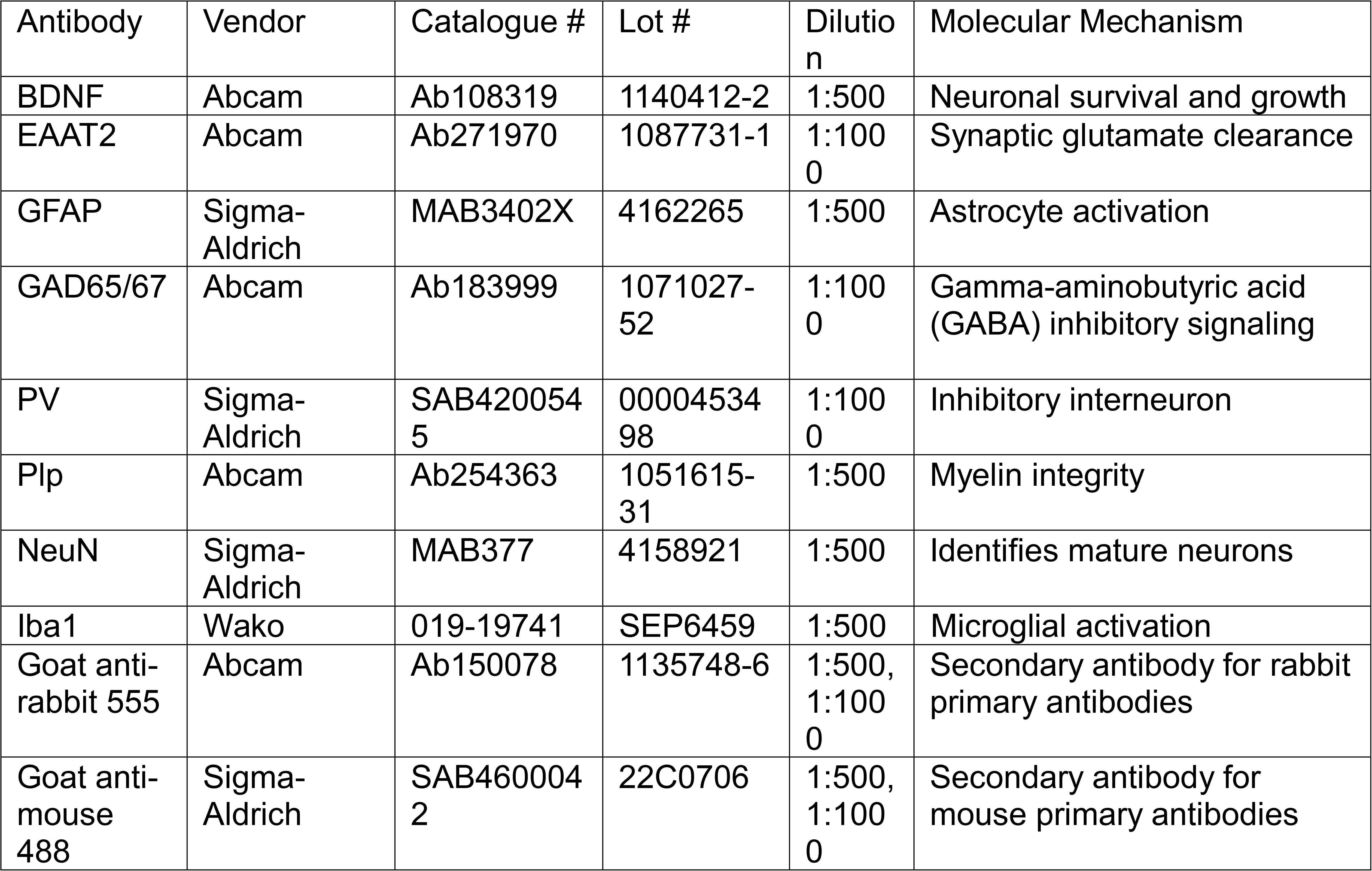
Antibodies and their molecular target used for immunofluorescence detection and analyses.

## REFERENCES

1. 2025 Alzheimer’s disease facts and figures. (2025). Alzheimer’s & Dementia, 21(4), e70235. 10.1002/alz.70235

2. Abdulai-Saiku, S., Gupta, S., Wang, D., Marino, F., Moreno, A. J., Huang, Y., Srivastava, D., Panning, B., & Dubal, D. B. (2025). The maternal X chromosome affects cognition and brain ageing in female mice. Nature, 638(8049), 152–159. 10.1038/s41586-024-08457-y

3. Anastasiades, P. G., & Carter, A. G. (2021). Circuit organization of the rodent medial prefrontal cortex. Trends in Neurosciences, 44(7), 550–563. 10.1016/j.tins.2021.03.006

4. Andrawus, M., Sharvit, L., & Atzmon, G. (2022). Epigenetics and Pregnancy: Conditional Snapshot or Rolling Event. International Journal of Molecular Sciences, 23(20), 12698. 10.3390/ijms232012698

5. Barba-Müller, E., Craddock, S., Carmona, S., & Hoekzema, E. (2019). Brain plasticity in pregnancy and the postpartum period: Links to maternal caregiving and mental health. Archives of Women’s Mental Health, 22(2), 289–299. 10.1007/s00737-018-0889-z

6. Basit, S., Wohlfahrt, J., & Boyd, H. A. (2023). Associations between parenthood and dementia in men and women: Biology or confounding? BMC Neurology, 23, 90. 10.1186/s12883-023-03108-7

7. Bathina, S., & Das, U. N. (2015). Brain-derived neurotrophic factor and its clinical implications. Archives of Medical Science : AMS, 11(6), 1164–1178. 10.5114/aoms.2015.56342

8. Beckman, M., Knox, K., Koneval, Z., Smith, C., Jayadev, S., & Barker-Haliski, M. (2020). Loss of presenilin 2 age-dependently alters susceptibility to acute seizures and kindling acquisition. Neurobiology of Disease, 136, 104719. 10.1016/j.nbd.2019.104719

9. Beeri, M. S., Rapp, M., Schmeidler, J., Reichenberg, A., Purohit, D. P., Perl, D. P., Grossman, H. T., Prohovnik, I., Haroutunian, V., & Silverman, J. M. (2009). Number of children is associated with neuropathology of Alzheimer’s disease in women. Neurobiology of Aging, 30(8), 1184–1191. 10.1016/j.neurobiolaging.2007.11.011

10. Belloy, M. E., Guen, Y. L., Stewart, I., Williams, K., Herz, J., Sherva, R., Zhang, R., Merritt, V., Panizzon, M. S., Hauger, R. L., Gaziano, J. M., Logue, M., Napolioni, V., & Greicius, M. D. (2024). Role of the X Chromosome in Alzheimer Disease Genetics. JAMA Neurology, 81(10), 1032–1042. 10.1001/jamaneurol.2024.2843

11. Bin, X., & Luo, H. (2025). Effects of pregnancy on neuron electrophysiology in prefrontal cortex, hippocampus, and basolateral amygdala of mouse brain. Animal Models and Experimental Medicine, 8(5), 864–873. 10.1002/ame2.12574

12. Bjerke, I. E., Yates, S. C., Laja, A., Witter, M. P., Puchades, M. A., Bjaalie, J. G., & Leergaard, T. B. (2020). Densities and numbers of calbindin and parvalbumin positive neurons across the rat and mouse brain. iScience, 24(1), 101906. 10.1016/j.isci.2020.101906

13. Blauwendraat, C., Wilke, C., Jansen, I. E., Schulte, C., Simón-Sánchez, J., Metzger, F. G., Bender, B., Gasser, T., Maetzler, W., Rizzu, P., Heutink, P., & Synofzik, M. (2016). Pilot whole-exome sequencing of a German early-onset Alzheimer’s disease cohort reveals a substantial frequency of *PSEN2* variants. Neurobiology of Aging, 37, 208.e11–208.e17. 10.1016/j.neurobiolaging.2015.09.016

14. Bourzac, K. (2025). Why women experience Alzheimer’s disease differently from men. Nature, 640(8059), S14–S17. 10.1038/d41586-025-01106-y

15. Bowers, G., Cullinan, W. E., & Herman, J. P. (1998). Region-Specific Regulation of Glutamic Acid Decarboxylase (GAD) mRNA Expression in Central Stress Circuits. The Journal of Neuroscience, 18(15), 5938–5947. 10.1523/JNEUROSCI.18-15-05938.1998

16. Cai, Y., An, S. S. A., & Kim, S. (2015). Mutations in presenilin 2 and its implications in Alzheimer’s disease and other dementia-associated disorders. Clinical Interventions in Aging, 10, 1163– 1172. 10.2147/CIA.S85808

17. Cho, W.-H., Noh, K., Lee, B. H., Barcelon, E., Jun, S. B., Park, H. Y., & Lee, S. J. (2022). Hippocampal astrocytes modulate anxiety-like behavior. Nature Communications, 13(1), 6536. 10.1038/s41467-022-34201-z

18. Colucci, M., Cammarata, S., Assini, A., Croce, R., Clerici, F., Novello, C., Mazzella, L., Dagnino, N., Mariani, C., & Tanganelli, P. (2006). The number of pregnancies is a risk factor for Alzheimer’s disease. European Journal of Neurology, 13(12), 1374–1377. 10.1111/j.1468-1331.2006.01520.x

19. Contador, J., Pérez-Millan, A., Guillén, N., Sarto, J., Tort-Merino, A., Balasa, M., Falgàs, N., Castellví, M., Borrego-Écija, S., Juncà-Parella, J., Bosch, B., Fernández-Villullas, G., Ramos-Campoy, O., Antonell, A., Bargalló, N., Sanchez-Valle, R., Sala-Llonch, R., & Lladó, A. (2022). Sex differences in early-onset Alzheimer’s disease. European Journal of Neurology, 29(12), 3623– 3632. 10.1111/ene.15531

20. Course, M. M., Gudsnuk, K., Keene, C. D., Bird, T. D., Jayadev, S., & Valdmanis, P. N. (2022). Aberrant splicing of PSEN2, but not PSEN1, in individuals with sporadic Alzheimer’s disease. Brain, 146(2), 507–518. 10.1093/brain/awac294

21. Cui, J., Jothishankar, B., He, P., Staufenbiel, M., Shen, Y., & Li, R. (2014). Amyloid precursor protein mutation disrupts reproductive experience-enhanced normal cognitive development in a mouse model of Alzheimer’s disease. Molecular Neurobiology, 49(1), 103–112. 10.1007/s12035-013-8503-x

22. Deacon, R. M. J., & Rawlins, J. N. P. (2006). T-maze alternation in the rodent. Nature Protocols, 1(1), 7–12. 10.1038/nprot.2006.2

23. Deems, N., & Leuner, B. (2020). Pregnancy, Postpartum and Parity: Resilience and Vulnerability in Brain Health and Disease. Frontiers in Neuroendocrinology, 57, 100820. 10.1016/j.yfrne.2020.100820

24. Duarte-Guterman, P., Richard, J. E., Lieblich, S. E., Eid, R. S., Lamers, Y., & Galea, L. A. M. (2023). Cellular and molecular signatures of motherhood in the adult and ageing rat brain. Open Biology, 13(11), 230217. 10.1098/rsob.230217

25. Eid, R. S., Chaiton, J. A., Lieblich, S. E., Bodnar, T. S., Weinberg, J., & Galea, L. A. M. (2019). Early and late effects of maternal experience on hippocampal neurogenesis, microglia, and the circulating cytokine milieu. Neurobiology of Aging, 78, 1–17. 10.1016/j.neurobiolaging.2019.01.021

26. Emrani, S., & Sundermann, E. E. (2025). Sex/gender differences in the clinical trajectory of Alzheimer’s disease: Insights into diagnosis and cognitive reserve. Frontiers in Neuroendocrinology, 77, 101184. 10.1016/j.yfrne.2025.101184

27. Ferguson, B. R., & Gao, W.-J. (2018). PV Interneurons: Critical Regulators of E/I Balance for Prefrontal Cortex-Dependent Behavior and Psychiatric Disorders. Frontiers in Neural Circuits, 12, 37. 10.3389/fncir.2018.00037

28. Fox, M., Berzuini, C., Knapp, L. A., & Glynn, L. M. (2018). Women’s Pregnancy Life History and Alzheimer’s Risk: Can Immunoregulation Explain the Link? American Journal of Alzheimer’s Disease & Other Dementias®, 33(8), 516–526. 10.1177/1533317518786447

29. Fox, M. M., Bramen, J. E., Kwon, D., Espeland, M. A., Sun, Y., Zhang, L., Shadyab, A. H., Casanova, R., Manson, J. E., Chang, M. Y., Porter, V. R., Farland, L. V., Xiao, Q., Wild, R. A., Siddarth, P., & Crandall, C. J. (2026). Pregnancy and breastfeeding are associated with less later-life cognitive decline in a longitudinal, prospective cohort. Alzheimer’s & Dementia: The Journal of the Alzheimer’s Association, 22(1), e71072. 10.1002/alz.71072

30. Gadek, M., Shaw, C. K., Abdulai-Saiku, S., Saloner, R., Marino, F., Wang, D., Bonham, L. W., Yokoyama, J. S., Panning, B., Benayoun, B. A., Casaletto, K. B., Ramani, V., & Dubal, D. B. (2025). Aging activates escape of the silent X chromosome in the female mouse hippocampus. Science Advances, 11(10), eads8169. 10.1126/sciadv.ads8169

31. Gilfarb, R. A., & Leuner, B. (2022). GABA System Modifications During Periods of Hormonal Flux Across the Female Lifespan. Frontiers in Behavioral Neuroscience, 16. 10.3389/fnbeh.2022.802530

32. Gong, J., Harris, K., Peters, S. A. E., & Woodward, M. (2022). Reproductive factors and the risk of incident dementia: A cohort study of UK Biobank participants. PLOS Medicine, 19(4), e1003955. 10.1371/journal.pmed.1003955

33. Gumusoglu, S. B., & Stelzer, I. A. (2025). Pregnancy as an opportunity to explore brain-immune connections in mental health. Science Advances, 11(10), eadu8270. 10.1126/sciadv.adu8270

34. Herreman, A., Hartmann, D., Annaert, W., Saftig, P., Craessaerts, K., Serneels, L., Umans, L., Schrijvers, V., Checler, F., Vanderstichele, H., Baekelandt, V., Dressel, R., Cupers, P., Huylebroeck, D., Zwijsen, A., Van Leuven, F., & De Strooper, B. (1999a). Presenilin 2 deficiency causes a mild pulmonary phenotype and no changes in amyloid precursor protein processing but enhances the embryonic lethal phenotype of presenilin 1 deficiency. Proceedings of the National Academy of Sciences of the United States of America, 96(21), 11872–11877.

35. Herreman, A., Hartmann, D., Annaert, W., Saftig, P., Craessaerts, K., Serneels, L., Umans, L., Schrijvers, V., Checler, F., Vanderstichele, H., Baekelandt, V., Dressel, R., Cupers, P., Huylebroeck, D., Zwijsen, A., Van Leuven, F., & De Strooper, B. (1999b). Presenilin 2 deficiency causes a mild pulmonary phenotype and no changes in amyloid precursor protein processing but enhances the embryonic lethal phenotype of presenilin 1 deficiency. Proceedings of the National Academy of Sciences, 96(21), 11872–11877. 10.1073/pnas.96.21.11872

36. Hol, E. M., & Pekny, M. (2015). Glial fibrillary acidic protein (GFAP) and the astrocyte intermediate filament system in diseases of the central nervous system. *Current Opinion in Cell Biology*, Cell Architecture, 32, 121–130. 10.1016/j.ceb.2015.02.004

37. Jang, H., Bae, J. B., Dardiotis, E., Scarmeas, N., Sachdev, P. S., Lipnicki, D. M., Han, J. W., Kim, T. H., Kwak, K. P., Kim, B. J., Kim, S. G., Kim, J. L., Moon, S. W., Park, J. H., Ryu, S.-H., Youn, J. C., Lee, D. Y., Lee, D. W., Lee, S. B., … Kim, K. W. (2018). Differential effects of completed and incomplete pregnancies on the risk of Alzheimer disease. Neurology, 91(7), e643–e651. 10.1212/WNL.0000000000006000

38. Jayadev, S., Case, A., Alajajian, B., Eastman, A. J., Möller, T., & Garden, G. A. (2013). Presenilin 2 influences miR146 level and activity in microglia. Journal of Neurochemistry, 127(5), 592–599. 10.1111/jnc.12400

39. Jayadev, S., Case, A., Eastman, A. J., Nguyen, H., Pollak, J., Wiley, J. C., Möller, T., Morrison, R. S., & Garden, G. A. (2010). Presenilin 2 Is the Predominant γ-Secretase in Microglia and Modulates Cytokine Release. PLoS ONE, 5(12), e15743. 10.1371/journal.pone.0015743

40. Jurisch-Yaksi, N., Sannerud, R., & Annaert, W. (2013). A fast growing spectrum of biological functions of γ-secretase in development and disease. Biochimica Et Biophysica Acta, 1828(12), 2815– 2827. 10.1016/j.bbamem.2013.04.016

41. Kapasi, A., DeCarli, C., & Schneider, J. A. (2017). Impact of Multiple Pathologies on the Threshold for Clinically Overt Dementia. Acta Neuropathologica, 134(2), 171–186. 10.1007/s00401-017-1717-7

42. Kim, D. Y., Ingano, L. A. M., Carey, B. W., Pettingell, W. H., & Kovacs, D. M. (2005). Presenilin/γ- Secretase-mediated Cleavage of the Voltage-gated Sodium Channel β2-Subunit Regulates Cell Adhesion and Migration*. Journal of Biological Chemistry, 280(24), 23251–23261. 10.1074/jbc.M412938200

43. Kinsley, C. H., Madonia, L., Gifford, G. W., Tureski, K., Griffin, G. R., Lowry, C., Williams, J., Collins, J., McLearie, H., & Lambert, K. G. (1999). Motherhood improves learning and memory. Nature, 402(6758), 137–138. 10.1038/45957

44. Knox, K. M., Beckman, M., Smith, C. L., Jayadev, S., & Barker-Haliski, M. (2023). Chronic seizures induce sex-specific cognitive deficits with loss of presenilin 2 function. Experimental Neurology, 361, 114321. 10.1016/j.expneurol.2023.114321

45. Ladyman, S. R., Khant Aung, Z., & Grattan, D. R. (2018). Impact of Pregnancy and Lactation on the Long-Term Regulation of Energy Balance in Female Mice. Endocrinology, 159(6), 2324–2336. 10.1210/en.2018-00057

46. Lee, B. H., Cevizci, M., Lieblich, S. E., Ibrahim, M., Wen, Y., Eid, R. S., Lamers, Y., Duarte-Guterman, P., & Galea, L. A. M. (2024). Exploring the parity paradox: Differential effects on neuroplasticity and inflammation by APOEe4 genotype at middle age. *Brain*, Behavior, and Immunity, 120, 54–70. 10.1016/j.bbi.2024.05.019

47. Lin, C.-L. G., Kong, Q., Cuny, G. D., & Glicksman, M. A. (2012). Glutamate transporter EAAT2: A new target for the treatment of neurodegenerative diseases. Future Medicinal Chemistry, 4(13), 1689–1700. 10.4155/fmc.12.122

48. Luquez, T., Algoo, J., Chiu, R., Mares, J. A., Yadav, A., Lam, M., Gaur, P., Lai, X., Lee, D. I., Paryani, F., Batchelor, R., Belli, I., Henry, J., Hoter-Ishay, B., Mattison, C., Starr, L., Lama, T., Karaahmet, B., Cao, W., … Menon, V. (2026). Cell-type signatures of Alzheimer’s disease shared across population groups. Nature, 657(8130), 191–201. 10.1038/s41586-026-10793-0

49. Macbeth, A. H., Gautreaux, C., & Luine, V. N. (2008). Pregnant rats show enhanced spatial memory, decreased anxiety, and altered levels of monoaminergic neurotransmitters. Brain Research, 1241, 136–147. 10.1016/j.brainres.2008.09.006

50. Martin, D. L., & Rimvall, K. (1993). Regulation of gamma-aminobutyric acid synthesis in the brain. Journal of Neurochemistry, 60(2), 395–407. 10.1111/j.1471-4159.1993.tb03165.x

51. Moscoso, A., Silva-Rodríguez, J., Aldrey, J. M., Cortés, J., Pías-Peleteiro, J. M., Ruibal, Á., Aguiar, P., & Alzheimer’s Disease Neuroimaging Initiative. (2022). 18F-florbetapir PET as a marker of myelin integrity across the Alzheimer’s disease spectrum. European Journal of Nuclear Medicine and Molecular Imaging, 49(4), 1242–1253. 10.1007/s00259-021-05493-y

52. Ohsawa, K., Imai, Y., Sasaki, Y., & Kohsaka, S. (2004). Microglia/macrophage-specific protein Iba1 binds to fimbrin and enhances its actin-bundling activity. Journal of Neurochemistry, 88(4), 844– 856. 10.1046/j.1471-4159.2003.02213.x

53. Page, C. E., Shepard, R., Heslin, K., & Coutellier, L. (2019). Prefrontal parvalbumin cells are sensitive to stress and mediate anxiety-related behaviors in female mice. Scientific Reports, 9(1), 19772. 10.1038/s41598-019-56424-9

54. Papuć, E., & Rejdak, K. (2018). The role of myelin damage in Alzheimer’s disease pathology. Archives of Medical Science : AMS, 16(2), 345–351. 10.5114/aoms.2018.76863

55. Pawluski, J. L., & Galea, L. A. M. (2006). Hippocampal morphology is differentially affected by reproductive experience in the mother. Journal of Neurobiology, 66(1), 71–81. 10.1002/neu.20194

56. Peeters, G., Katelekha, K., Lawlor, B., & Demnitz, N. (2022). Sex differences in the incidence and prevalence of young-onset Alzheimer’s disease: A meta-analysis. International Journal of Geriatric Psychiatry, 37(1), 10.1002/gps.5612. 10.1002/gps.5612

57. Pike, C. J. (2017). Sex and the development of Alzheimer’s disease. Journal of Neuroscience Research, 95(1–2), 671–680. 10.1002/jnr.23827

58. Pizzo, P., Basso, E., Filadi, R., Greotti, E., Leparulo, A., Pendin, D., Redolfi, N., Rossini, M., Vajente, N., Pozzan, T., & Fasolato, C. (2020). Presenilin-2 and Calcium Handling: Molecules, Organelles, Cells and Brain Networks. Cells, 9(10), 2166. 10.3390/cells9102166

59. *PLP1 proteolipid protein 1 [Homo sapiens (human)]—Gene—NCBI*. (n.d.). Retrieved May 9, 2026, from https://www.ncbi.nlm.nih.gov/gene/5354

60. Ragan, C. M., Ahmed, E. I., Vitale, E. M., Linning-Duffy, K., Miller-Smith, S. M., Maguire, J., & Lonstein, J. S. (2022). Postpartum State, but Not Maternal Caregiving or Level of Anxiety, Increases Medial Prefrontal Cortex GAD65 and vGAT in Female Rats. Frontiers in Global Women’s Health, 2. 10.3389/fgwh.2021.746518

61. Ravndal, A., Fjell, A. M., Vidal-Piñeiro, D., Sørensen, Ø., Falch, E. S., Kropiunig, J., Garrido, P. F., Roe, J. M., Alatorre-Warren, J.-L., Sneve, M. H., Bartrés-Faz, D., Pascual-Leone, A., Brandmaier, A. M., Düzel, S., Kühn, S., Lindenberger, U., Nyberg, L., Watne, L. O., Henson, R. N., … Grydeland, H. (2025). Sex differences in healthy brain aging are unlikely to explain higher Alzheimer’s disease prevalence in women. Proceedings of the National Academy of Sciences, 122(42), e2510486122. 10.1073/pnas.2510486122

62. Rim, C., You, M.-J., Shin, D., Kim, H.-J., Sung, S., Kim, Y., Park, J., & Kwon, M.-S. (2026). Postpartum Brain Reveals Repopulating Process and Long-Lasting Proteomic Changes of Microglia. Glia, 74(1), e70088. 10.1002/glia.70088

63. Robinson-Cooper, L., Davidson, S., Koutoubi, R., Zhang, K., Park, H., & Barker-Haliski, M. (2026). Loss of presenilin 2 function age-dependently increases susceptibility to kainate-induced acute seizures and blunts hippocampal kainate-type glutamate receptor expression. Experimental Neurology, 397, 115586. 10.1016/j.expneurol.2025.115586

64. Rodrı guez, C., Guillamón, A., Pinos, H., & Collado, P. (2004). Postpartum changes in the GABAergic system in the bed nucleus of the accessory olfactory tract. Neurochemistry International, 44(3), 179–183. 10.1016/S0197-0186(03)00115-3

65. Seibenhener, M. L., & Wooten, M. C. (2015). Use of the Open Field Maze to Measure Locomotor and Anxiety-like Behavior in Mice. Journal of Visualized Experiments : JoVE, (96), 52434. 10.3791/52434

66. Sert, N. P. du, Hurst, V., Ahluwalia, A., Alam, S., Avey, M. T., Baker, M., Browne, W. J., Clark, A., Cuthill, I. C., Dirnagl, U., Emerson, M., Garner, P., Holgate, S. T., Howells, D. W., Karp, N. A., Lazic, S. E., Lidster, K., MacCallum, C. J., Macleod, M., … Würbel, H. (2020). The ARRIVE guidelines 2.0: Updated guidelines for reporting animal research. PLOS Biology, 18(7), e3000410. 10.1371/journal.pbio.3000410

67. Shea, Y.-F., Chu, L.-W., Chan, A. O.-K., Ha, J., Li, Y., & Song, Y.-Q. (2016). A systematic review of familial Alzheimer’s disease: Differences in presentation of clinical features among three mutated genes and potential ethnic differences. Journal of the Formosan Medical Association = Taiwan Yi Zhi, 115(2), 67–75. 10.1016/j.jfma.2015.08.004

68. Sheppard, O., & Coleman, M. (2020). Alzheimer’s Disease: Etiology, Neuropathology and Pathogenesis. In X. Huang (Ed.), Alzheimer’s Disease: Drug Discovery. Exon Publications. http://www.ncbi.nlm.nih.gov/books/NBK566126/

69. Sobow, T., & Kloszewska, I. K. (2003). Modulation of Age at Onset in Late-Onset Sporadic Alzheimer s Disease by Estrogen-Related Factors:The Age of Menopause and Number of Pregnancies. German Journal of Psychiatry, 6.

70. Tan, Y.-L., Yuan, Y., & Tian, L. (2020). Microglial regional heterogeneity and its role in the brain. Molecular Psychiatry, 25(2), 351–367. 10.1038/s41380-019-0609-8

71. Trainer, R., Stafisso-Sandoz, G., Quadros, P., Marcus, L. K., Hearon, C., Meyer, E. A. A., Hester, N., Morgan, M., Kozub, F. J., & Lambert, K. G. (2006). Motherhood and the hormones of pregnancy modify concentrations of hippocampal neuronal dendritic spines. Hormones and Behavior, 49(2), 131–142. 10.1016/j.yhbeh.2005.05.017

72. Wang, Y., Fu, A. K. Y., & Ip, N. Y. (2022). Instructive roles of astrocytes in hippocampal synaptic plasticity: Neuronal activity-dependent regulatory mechanisms. The Febs Journal, 289(8), 2202–2218. 10.1111/febs.15878

73. Xu, J., Kobayashi, S., Yamaguchi, S., Iijima, K., Okada, K., & Yamashita, K. (2000). Gender Effects on Age-Related Changes in Brain Structure. AJNR: American Journal of Neuroradiology, 21(1), 112–118.

74. Yan, L., Li, L., Han, W., Pan, B., Xue, X., & Mei, B. (2013). Age-related neuropsychiatric symptoms in *presenilins* conditional double knockout mice. Brain Research Bulletin, 97, 104–111. 10.1016/j.brainresbull.2013.06.002

75. Zhao, C., Driessen, T., & Gammie, S. C. (2012). Glutamic acid decarboxylase 65 and 67 expression in the lateral septum is up-regulated in association with the postpartum period in mice. Brain Research, 1470, 35–44. 10.1016/j.brainres.2012.06.002

76. Zhao, C., & Gammie, S. C. (2014). Glutamate, GABA, and glutamine are synchronously upregulated in the mouse lateral septum during the postpartum period. Brain Research, 1591, 53–62. 10.1016/j.brainres.2014.10.023

77. Zimberknopf, E., Xavier, G. F., Kinsley, C. H., & Felicio, L. F. (2011). Prior Parity Positively Regulates Learning and Memory in Young and Middle-Aged Rats. Comparative Medicine, 61(4), 366–377.

