## Supplemental Table 1 for "Pregnancy History and Loss of Normal PSEN2 Function Impact Female Late-Life Cognitive Processing, Inhibitory Signaling, and Gliosis"

**Fang et al.**

**Supplemental Table 1. Antibodies and their molecular target used for immunofluorescence detection and analyses.**

| Antibody | Vendor | Catalogue # | Lot # | Dilution | Molecular Mechanism |
| --- | --- | --- | --- | --- | --- |
| BDNF | Abcam | Ab108319 | 1140412-2 | 1:500 | Neuronal survival and growth |
| EAAT2 | Abcam | Ab271970 | 1087731-1 | 1:1000 | Synaptic glutamate clearance |
| GFAP | Sigma-Aldrich | MAB3402X | 4162265 | 1:500 | Astrocyte activation |
| GAD65/67 | Abcam | Ab183999 | 1071027-52 | 1:1000 | Gamma-aminobutyric acid (GABA) inhibitory signaling |
| PV | Sigma-Aldrich | SAB4200545 | 0000453498 | 1:1000 | Inhibitory interneuron |
| Plp | Abcam | Ab254363 | 1051615-31 | 1:500 | Myelin integrity |
| NeuN | Sigma-Aldrich | MAB377 | 4158921 | 1:500 | Identifies mature neurons |
| Iba1 | Wako | 019-19741 | SEP6459 | 1:500 | Microglial activation |
| Goat anti-rabbit 555 | Abcam | Ab150078 | 1135748-6 | 1:500, 1:1000 | Secondary antibody for rabbit primary antibodies |
| Goat anti-mouse 488 | Sigma-Aldrich | SAB4600042 | 22C0706 | 1:500, 1:1000 | Secondary antibody for mouse primary antibodies |
